# Probing cellular response to topography in three dimensions

**DOI:** 10.1101/232066

**Authors:** Colin D. Paul, Alex Hruska, Jack R. Staunton, Hannah A. Burr, Kathryn M. Daly, Jiyun Kim, Nancy Jiang, Kandice Tanner

**Affiliations:** Laboratory of Cell Biology, Center for Cancer Research, National Cancer Institute, National Institutes of Health

**Keywords:** Topographical cues, engineered matrices, microrheology, physical properties, cell alignment, cell protrusions

## Abstract

Biophysical aspects of in vivo tissue microenvironments include microscale mechanical properties, fibrillar alignment, and architecture or topography of the extracellular matrix (ECM). These aspects act in concert with chemical signals from a myriad of diverse ECM proteins to provide cues that drive cellular responses. Here, we used a bottom-up approach to build fibrillar architecture into 3D amorphous hydrogels using magnetic-field driven assembly of paramagnetic colloidal particles functionalized with three types of human ECM proteins found in vivo. We investigated if cells cultured in matrices comprised of fibrils of the same size and arranged in similar geometries will show similar behavior for each of the ECM proteins tested. We were able to resolve spatial heterogeneities in microscale mechanical properties near aligned fibers that were not observed in bulk tissue mechanics. We then used this platform to examine factors contributing to cell alignment in response to topographical cues in 3D laminin-rich matrices. Multiple human cell lines extended protrusions preferentially in directions parallel or perpendicular to aligned fibers independently of the ECM coating. Focal adhesion proteins, as measured by paxillin localization, were mainly diffuse in the cytoplasm, with few puncta localized at the protrusions. Integrin β1 and fascin regulated protrusion extension but not protrusion alignment. Myosin II inhibition did not reduce observed protrusion length. Instead, cells with reduced myosin II activity generated protrusions in random orientations when cultured in hydrogels with aligned fibers. Similarly, myosin II dependence was observed in vivo, where cells no longer aligned along the abluminal surfaces of blood vessels upon treatment with blebbistatin. These data suggest that myosin II can regulate sensing of topography in 3D engineered matrices for both normal and transformed cells.

## INTRODUCTION

The physical properties of the extracellular matrix (ECM) milieu are widely acknowledged as fundamental determinants of cell fate, tissue homeostasis, immune response, wound healing, and cancer progression [1–4]. Within a given tissue, the ECM not only provides structural support but regulates cell signaling via reciprocal biochemical and biophysical cues [5]. On one hand, the ECM contributes to overall tissue mechanics, and the effects of mechanical properties on cell fate and phenotype have been extensively studied using in vitro and in vivo assays [6–15]. For example, tissues become progressively stiffer as a function of malignant transformation from normal to tumors [9]. In addition, the architecture of the ECM provides structural feedback, such as topographical cues [13–16]. Topography, simply described, refers to the shape and profile of a given material’s surface [17–19]. Cells respond to topographical and stiffness-mediated cues through biochemical signaling via cellular adhesions and cytoskeletal attachments to the ECM [20]. Cell sensing of architectural and mechanical cues is a complex phenomenon where ECM adhesion molecules act concomitantly with intracellular machinery to drive cellular responses [21]. In vivo, tissue topographical cues are heterogeneous, with hybrid structures comprised of aligned ridges and pores that span lengths from nanoscale to microscale [17–19]. On the nanoscale level, ECM proteins can adopt different morphologies, such as globular and fibrillar architectures [17, 18]. One such example is fibronectin (FN), which presents different cell binding sites and is alternatively spliced to generate conformation which initiate distinct signaling cascades [22, 23]. On the other hand, the chemical specificity of these building blocks in turn also regulate unique signaling cascades. For example, cells that interact with fibronectin fibrils receive distinct chemical cues from those received when exposed to cues derived from collagen type I fibrils. Yet, in some cases, physical cues may dominate cellular response in the presence of a chemical cue. Simulations of stretching of a module of a FN fibril, FN III, is sufficient to override beta one-dependent modulation of increased ligand binding and associated down-stream signaling. Thus, understanding how cells “sense” these interconnected cues and how they influence eventual cell fate remains a perplexing issue.

These concepts are often difficult to discern using naturally derived 3D tissue mimetics, as precise control of ligand density and architecture are often intertwined. Moreover, dissecting differential physical cues such as mechanics from architecture is also challenging. While studies using patterned two-dimensional substrates [24, 25] and microfabricated environments [26] have revealed aspects of cell response to topography, reactions to topographic cues in more physiologically relevant three-dimensional environments are not as well understood. Importantly, three-dimensional cues are likely vital to recapitulate some aspects of physiological mechanosensing. Also, cell attachments to matrices in 3D environments often differ with respect to that observed for cells cultured in 2D substrates, which in turn may impact mechanosensing in specific 3D ECMs. In tissue, 3D topographies consist of highly oriented structures that are not well-recapitulated by *in vitro* hydrogel models. For example, commonly used collagen hydrogels form fibers that are randomly oriented unless some external micropatterning is imposed during their polymerization [27]. Similarly, laminin-rich ECMs (Matrigel) form amorphous gels devoid of cell-scale structures [28]. To address the need for reproducible 3D culture systems with well-defined matrix architecture and ECM protein composition, we recently developed a method whereby functionalized paramagnetic colloidal particles are magnetically aligned in 3D hydrogels to create fibrils that span microns in length and 10s of nms in widths [17, 29]. Fiber alignment, diameter, spacing, and extracellular matrix conjugation to the colloidal particles can be controlled to create defined topography independently of the ligand used to coat the particles. In 3D Matrigel matrices containing aligned particles, mouse fibroblasts and neural cell lines send out protrusions that are longer than those seen in matrices lacking particle alignment, independently of the ECM ligand conjugated to the colloidal particles. When the particles are coated in fibronectin, these cells preferentially extend protrusions either parallel or perpendicular to the fibers. This system allows us to test if cells cultured in matrices where fibrils of the same size and arranged in similar geometries will show similar behavior for different types of ECM proteins. In this system, the bulk mechanical properties were similar regardless of the presence of aligned or unaligned colloidal particles. However, lack of understanding of mechanical cues at the cellular scale, of how ECM nanoparticle conjugation affects cell response, and of mechanistic factors driving cell response to topography limited the use of this system.

Here, we used our system to discern the role of aligned topographical cues, presented across a range of human ECM proteins, on human cell response in Matrigel (laminin-rich) matrices with well-characterized physical properties. Using optical trap-based active microrheology, we measured the 3D microscale viscoelasticity. We then asked how topographical and micromechanical cues influence human normal (human foreskin fibroblast, HFF) and cancer (U87 glioblastoma) cells on this length scale. Using genetic manipulation and small molecule inhibitors, we determined that β1integrin and fascin reduced the length of cell protrusions in response to physical cues resulting from fiber alignment in these engineered Matrigel matrices. However, protrusions were still aligned with the fibrils. In contrast, reduced myosin II activity did not affect protrusion length, but protrusions were randomly oriented. We confirmed that myosin II is also required by cells to sense topographical alignment in vivo using the zebrafish brain vasculature as our model system. Our results suggest that normal and cancer cells use similar machinery to respond to the topographical and micromechanical cues in this system, where myosin II may regulate how cells sense topographical cues.

## MATERIALS AND METHODS

### Cell culture

Human foreskin fibroblast cells were cultured in Dulbecco’s Modified Eagles Medium (DMEM) supplemented with 10% fetal bovine serum, 1% penicillin-streptomycin, and 1% L-glutamine and maintained at 37°C and 10% CO2. Human U87 glioma cells were cultured in the same medium but maintained at 37°C and 5% CO2. Cells were sub-passaged every 2-3 days.

### Conjugation of fluorophores to human proteins

We employed several proteins. Human laminin (pepsinized; Millipore Sigma, Burlington, MA, Catalog #AG56P), human tenascin-C (Millipore Sigma, Catalog #CC065), and human plasma fibronectin (Millipore Sigma, Catalog #FC010) were conjugated to fluorophores using the DyLight™ 488 Microscale Labeling Kit (ThermoFisher Scientific, Waltham, MA, Catalog #53025) according to the manufacturer’s instructions. Briefly, proteins were supplied in suspension at concentrations ranging from 0.25-1 mg/ml. Laminin and tenascin-C were at concentrations less than recommended for labeling (0.5 mg/ml and 0.25 mg/ml, respectively; suggested concentration for conjugation to fluorophore is 1 mg/ml). For these proteins, 50 μg and 25 μg, respectively, were used in the labeling reaction instead of the recommended 100 μg. Bovine serum albumin (BSA) dissolved in PBS at a concentration of 1 mg/ml was also labeled for use as a control that does not specifically bind to cell adhesion proteins. Following labeling, protein concentrations were measured using a DeNovix DS-11+ Spectrophotometer and the “Labeled Protein” module. E1% was set at 10 g/100 ml, 1A = 1 mg/ml, with analysis wavelength for fluorescence at 494 nm, extinction coefficient set at 71,000, A260 factor of 0.3, and A280 factor of 0.11. Successful labeled was indicated by an absorption peak at 494 nm.

### Conjugation of proteins to magnetic colloidal particles

Labeled proteins were conjugated to 300-nm diameter paramagnetic colloidal particles as described previously using the Ademtech Carboxy-Adembeads Coupling Kit (Ademtech, Pessac, France, Catalog #02820) [17]. Briefly, 0.5 mg of carboxylated superparamagnetic colloidal particles were washed twice in 100 μl Activation Buffer before being resuspended in 100 μl of fresh Activation Buffer. The beads were then activated by the addition of 100 μl of 4 mg/ml EDC (resuspended in Activation Buffer) to the colloidal particles in solution, and this mixture was incubated for 1 h at room temperature. To conjugate proteins to the beads, 20 μg of fluorescently-labeled protein in solution was added to the tube, and the mixture was incubated overnight under gentle shaking. Then, 200 μl of 0.5 mg/ml BSA in Activation Buffer was added to the tube and incubated while shaking for 1 h at room temperature to quench the reaction. Functionalized beads were washed three times with 100 μl Storage Buffer, and beads were then stored at a final concentration of 10 mg/ml in Storage Buffer at 4°C for up to 2 weeks.

### Cell seeding and topographic alignment in 3D matrices

Matrigel (BD Corning, Corning, NY, Catalog #356230) was thawed on ice and maintained at 4°C. Prior to experiments, Matrigel was stored on ice to reach a stable temperature. A 100 μl aliquot of Matrigel was spread uniformly over the surface of a Lak-Tek 4-well chambered coverglass well (ThermoFisher, Catalog #155383) chilled on ice and placed in an incubator at 37°C for 5 minutes to polymerize. Prior to seeding in 3D culture, HFF or U87 cells were washed with phosphate buffered saline (PBS), detached from the cell culture flask using 10 mM EDTA in PBS, washed once in growth medium, centrifuged at 1000 rpm for 5 min, and resuspended to a concentration of 2 × 106 cells/ml in serum-free medium. Cells were then mixed with functionalized colloidal particles and Matrigel in the following ratio: 12 μl beads at a concentration of 10 mg/ml, 430 μl Matrigel, and 50 μl cell suspension (100,000 cells).

The mixture was gently mixed by pipette and added to a well of the 4-well coverslide. For alignment of the colloidal particles, the slide was immediately placed on a magnet (NdFeB magnet, K&J Magnetics, Pipersville, PA, Catalog #BX8X8X8, 25.4 mm x 25.4 mm x 25.4 mm) chilled in ice for 15 minutes. Following alignment, the slide was placed in the 37°C incubator for 30 minutes to polymerize the Matrigel. For unaligned topographies, slides were placed on ice far from the magnet for 15 minutes following the addition of the cell/nanoparticle/Matrigel mixture and then polymerized for 30 minutes at 37°C. Wells on the slides were seeded sequentially to ensure proper nanoparticle alignment over the center of the magnet, where field lines are parallel. Following Matrigel polymerization, 350 μl of serum-free medium was added to each well. Slides were placed in the incubator for 24 h prior to fixation. As a control, an equal number of cells was seeded in a Matrigel matrix, where the 12 μl bead mixture was replaced with pure Matrigel, resulting in a matrix containing cells but no additional ECM-conjugated colloidal particles.

To seed cells in aligned and unaligned agarose matrices, a solution of 1% agarose (Millipore Sigma, Catalog #A9414-25G) dissolved in serum free media and sterilized with a 0.22 μm filter was prepared. Chamber slides were coated with 300 μl/well agarose. The cell solution, particles, and warmed agarose were mixed in the volumes indicated above, and particles were placed on the magnet for alignment (or kept away from the magnet for unaligned matrices) in a 40°C incubator for 10 minutes. Slides were then kept at room temperature for 10 minutes for gels to set. A volume of 350 μl of serum free media was added to the gels, and gels were kept in the incubator for 48 hours prior to fixation and imaging.

Hyaluronic acid matrices were prepared using the HyStem Cell Culture Scaffold Kit (Millipore Sigma, Catalog #HYS020). HyStem, Extralink 1, and degassed water were allowed to come to room temperature. Under aseptic conditions, using a syringe and needle, 1.0 ml of degassed water was added to the HyStem bottle containing 10 mg of HyStem, and 0.5 ml degassed water was added to the Extralink 1 bottle containing 5 mg of Extralink. To form the hydrogel, a ratio of one part Extralink to three parts HyStem was used. A 100 μl aliquot of hydrogel was spread uniformly over the surface of a chamber slide and allowed to polymerize at room temperature for 15 minutes. Cells (prepared as described above) were mixed with functionalized nanoparticles and hydrogel in the following ratio: 12 μl beads at a concentration of 10 mg/ml, 430 μl HyStem at concentration of 10 mg/ml, 143 μl of Extralink 1 at a concentration of 0.5 mg/ml, and 50 μl cell suspension (100,000 cells). The mixture was gently mixed by pipette, and 635 μl of this mixture was added to a well of the 4-well coverslide. For alignment of the nanoparticles, the slide was immediately placed on the magnet at room temperature for 15 minutes. For unaligned topographies, slides were placed far from the magnet for 15 minutes. Following alignment, the slide was allowed to fully polymerize in the 37°C incubator for 30 minutes. Following hydrogel polymerization, 350 μl of serum-free medium was added to each well. After 24 h, media was replaced with serum-containing media (10% FBS). Cells were fixed and analyzed 24 h after the media change.

### Pharmacological inhibition

Cell contractility was assessed using a myosin II inhibitor (blebbistatin) compared to a vehicle control. For blebbistatin experiments, serum-free media was supplemented with 100 μM (-)-blebbistatin (Millipore Sigma, Catalog #B0560) or vehicle control (VC; DMSO), and 350 μl of media was added to embedded cells after matrix topography was set as described above and prior to overnight incubation. Inhibition of fascin was obtained by use of fascin-G2 (Xcessbio, San Diego, CA, Catalog #M60269-2s), where serum-free media was supplemented with 125 μM of the inhibitor or vehicle control (VC; DMSO), and 350 μl of media was added to embedded cells after matrix topography was set as described above and prior to overnight incubation. In these experiments, matrices containing fibronectin--conjugated colloidal particles were used. For cells blocked with a function-blocking antibody, cells were resuspended at a concentration of 2×10^6^ cells/ml in serum-free media containing 30 μg/ml of an IgG mouse isotype control antibody (abcam, Cambridge, MA, Catalog #ab91353) or a function blocking anti-β1 integrin antibody (abcam, Catalog #ab24693). Cells were seeded in aligned matrices with fibronectin-conjugated colloidal particles as described above. After matrix topography was set, 350 μl of serum-free media supplemented with 30 μg/ml of isotype control or function-blocking antibody was added to each well, and the sample was incubated overnight.

### Knockdown of β1 integrin via siRNA

To knockdown expression of β1 integrin, cells were detached with 0.25% trypsin-EDTA and plated at 200,000 cells/well in a 6-well plate in 2.5 ml total growth medium. The following day, cells were transfected with ThermoFisher Silencer Select siRNA targeting expression of β1 integrin (ThermoFisher, Catalog #4390824, siRNA ID s7575), or with a negative control (Silencer Select Negative Control No. 1, ThermoFisher, Catalog #4390843). siRNA stored at a concentration of 50 μM and stored at −20°C and diluted 1:10 in Opti-MEM cell culture medium (ThermoFisher, Catalog #31985070) to reach a concentration of 5 μM prior to transfection. Per well of a 6-well plate, 3 μl of siRNA at 5 μM was mixed with 150 μl Opti-MEM. In a separate tube, 7.2 μl RNAiMAX (ThermoFisher, Catalog #13778030) was mixed with 150 μl Opti-MEM. 150 μl diluted of the Opti-MEM/siRNA mixture was combined with 150 μl of the Opti-MEM/RNAiMAX mixture and incubated for 5 min at room temperature. To transfect, 250 μl of this mixture was added to cells plated the previous day, directly to the existing medium. The following day, media was replaced with fresh growth media. After 48 h, cells were prepared for seeding in aligned matrices as described above.

To assess knockdown, RNA was isolated from one representative biological replicate of transfected cells using Trizol (ThermoFisher, Catalog #15596018) according to the manufacturer’s instructions. Briefly, 0.4 ml Trizol was added directly to cells plated in 6-well plate wells after aspirating media. The lysate was pipetted up and down to homogenize, mixed with 0.2 ml chloroform per ml of Trizol reagent, and centrifuged for 15 minutes at 12,000 × g at 4 °C. The resulting upper aqueous phase containing RNA was mixed with an equal volume of 100% ethanol and processed with the PureLink RNA Mini Kit (ThermoFisher, Catalog #12183020). The mixture was processed through the kit Spin Cartridge by centrifugation at 12,000 × g for 1 minute, washed once with 700 μl per tube of Wash Buffer I, washed twice with 500 μl per tube of Wash Buffer II, and eluted in RNase-free water. The resulting RNA concentration was assessed via Nanodrop.

cDNA was synthesized using SuperScript IV VILO Master Mix (ThermoFisher, Catalog #11755050). For each sample, 10 μl of RNA at 100 ng/μl was combined with 4 μl of SuperScript IV VILO Master Mix or No RT Control, and 6 μl nuclease-free water was added to bring the total volume to 20 μl. Solutions were gently mixed and incubated at 25°C for 10 minutes, 50°C for 10 minutes, and 85°C for 5 minutes.

The synthesized cDNA was used for RT-PCR. For each well of a 96-well reaction plate, 10 μl TaqMan Fast Advanced Master Mix (ThermoFisher, Catalog #4444557) was mixed with 2 μl of the synthesized cDNA, 7 μl nuclease-free water, and 1 μl of the TaqMan probe. Samples were run in technical triplicates. The FAM-MGB TaqMan probes used were: GAPDH (ThermoFisher, Catalog #4331182, Assay ID Hs04420697_g1) and ITGB1 (ThermoFisher, Catalog #4331182, Assay ID Hs01127536_m1).

Gene expression assays were performed on an Applied Biosystems QuantStudio 3 (ThermoFisher). Gene expression was normalized to that of GAPDH for each cell type and siRNA status. For cells transfected with siRNA targeting integrin β1, fold change was calculated from the mean ∆∆cT of three technical triplicates, as indicated by the manufacturer (Applied Biosystems, “Guide to Performing Relative Quantitation of Gene Expression Using Real-Time Quantitative PCR”). For cells transfected with control siRNA, expression of integrin β1 was set to 1.

### Transduction of paxillin and actin biosensors

HFF cells were plated in 24-well plates the day prior to transduction, with 50,000 cells/well in 1 ml of growth medium. Cells were transduced with a LentiBrite™ paxillin-GFP lentiviral biosensor (Millipore Sigma, Catalog #17-10154) at a MOI of 20 in 1 ml of growth media per well. Transduction media contained 5 μg/ml polybrene transfection reagent (Millipore Sigma, Catalog #TR-1003-G). After 24 h, transduction media was replaced with fresh growth media. After an additional 24 h incubation, cells were plated in 3D matrices as described above. Cells were concurrently plated on 2D surfaces coated with Matrigel to assess GFP-paxillin expression on surfaces where focal adhesions were expected to form.

Similarly, U87 cells were transfected with a LifeAct TagRFP adenoviral vector (rAV-CMV-LifeAct, ibidi, Martinsried, Germany, Catalog #60122). Briefly, 1 million U87 cells were seeded in a T75 flask and infected at a MOI of 15 in 5 ml serum free media containing 26.6 μl ibiBoost adenovirus transduction enhancer (ibidi, Catalog #50301). Media containing virus was removed and replaced with fresh media after 4 h. Live cells expressing LifeAct were imaged immediately after seeding in 3D Matrigel matrices containing fibronectin-conjugated colloidal particles.

### Cell fixation and fluorescent staining

For samples embedded in Matrigel matrices (with or without colloidal particles), media was aspirated, and each well was washed with 200 μl PBS. To each well, 300 μl of 4% paraformaldehyde diluted in PBS was added, and samples were incubated for 3 hours at room temperature. Fixed samples were washed three times with 200 μl/well PBS. Samples were then incubated with 300 μl/well of 1% BSA in PBS at room temperature for 30 minutes to block non-specific binding and subsequently washed twice with 200 μl/well PBS. A phalloidin stock solution was prepared by dissolving 10 nmol of phalloidin-Atto 565 (Millipore Sigma, Catalog #94072-10NMOL) in 500 μl methanol. To fluorescently label F-actin and the nucleus, a solution containing 2 μg/ml Hoechst 33258 (ThermoFisher, Catalog #H3569) and 30 μl phalloidin stock/ml in 1% BSA in PBS was prepared. Cells were stained with 200 μl/well of the staining solution at room temperature for 3 h, then washed three times with 200 μl/well PBS. Samples were stored in PBS at 4°C prior to imaging. Cells transduced with the GFP-paxillin biosensor were prepared following the same procedure, but the actin cytoskeleton was stained with a 1:40 dilution of AlexaFluor™ 633 phalloidin (ThermoFisher, Catalog #A22284).

To visualize focal adhesion formation in 2D environments in the absence of transduction with the GFP-paxillin biosensor, HFF cells were plated in ibidi μ-Slide VI0.4 channel slides with ibiTreat surfaces (ibidi, Catalog #80606). After attaching in the presence of serum, cells were fixed for 15 min at room temperature using 4% paraformaldehyde diluted in PBS. Channels were washed three times with PBS, and non-specific binding was blocked for 1 h at room temperature using a solution of 0.3% Triton X-100 and 20% goat serum in PBS. After blocking, channels were washed three times with PBS. An anti-paxillin antibody (purified mouse anti-paxillin, close 349, BD Biosciences, San Jose, CA, Catalog #610052) was diluted 1:75 in PBS containing 1% BSA, added to the cells, and incubated at 4°C overnight. Following incubation, cells were washed three times in PBS. A secondary antibody solution containing a 1:200 dilution of goat anti-mouse AlexaFluor™ 594 (ThermoFisher, Catalog# A11020), a 1:40 dilution of AlexaFluor™ 488 phalloidin (ThermoFisher, Catalog #A12379), and 1 μg/ml Hoechst 33342 (ThermoFisher, Catalog #H3570) diluted in 1% BSA in PBS was added and incubated at room temperature for 1 h. Cells were washed five times with PBS prior to imaging.

### Confocal microscopy

For cells embedded in 3D matrices, images were acquired on a Zeiss 780 LSM confocal microscope. One-photon, confocal, 12-bit, 2-dimensional images were acquired at lateral dimensions of 512×512 pixels with a Zeiss 20× Plan-Apochromat, 0.8 NA objective. Individual images were tiled (3×3 grid) to image a total area of 1275.29 μm × 1275.29 μm (1536 pixels × 1536 pixels). Tiled images were acquired in z-stacks spaced 2 μm apart over an axial distance of ~120 μm to image cells throughout the matrix. Samples were excited with 561 nm light from a solid-state laser with a total power of 20 mW, 405 nm light from a laser diode with a total power of 30 mW, and 488 nm light from an argon laser with a total power of 25 mW. Lasers were set at or below 2.4% of the total power. Two beam splitters, MBS 488/561 and MBS 405, were employed in the emission pathway to delineate the red, green, and blue channels. Transmitted light was also collected. Pinhole width was set at 90 μm. Pixel dwell was set at 1.58 μs. The master gain was set at or below 890 for all images acquired. For some images in Figures 1-3, confocal z stacks were acquired at 2048 pixels x 2048 pixels to obtain greater detail of cell and fiber morphology. To image expression of paxillin in 3D matrices, one-photon, confocal, 12-bit, 2-dimensional images were acquired at lateral dimensions of 512×512 pixels with a Zeiss 40× Plan-Apochromat, 1.4 NA oil immersion objective on a Zeiss 780 LSM confocal microscope. Images were acquired at 4 times digital zoom to achieve a final pixel size of 0.1038 μm × 0.1038 μm. Images were acquired in z-stacks spaced 1 μm apart over the height of a cell. Samples were excited with 633 nm light from a solid-state laser with a total power of 5 mW, 405 nm light from a laser diode with a total power of 30 mW, and 488 nm light from an argon laser with a total power of 25 mW. Lasers were set at or below 4% of the total power. Tracks were imaged in sequence to minimize crosstalk. Pinhole width was set at 41.3 μm. Pixel dwell was set at 1.58 μs. The master gain was set at or below 700 for all images acquired. Axial stacks were trimmed to contain only the images with the fibrils in focus, and maximum intensity projections were made to visualize the actin cytoskeleton, nucleus, and paxillin expression.

**Figure 1.**
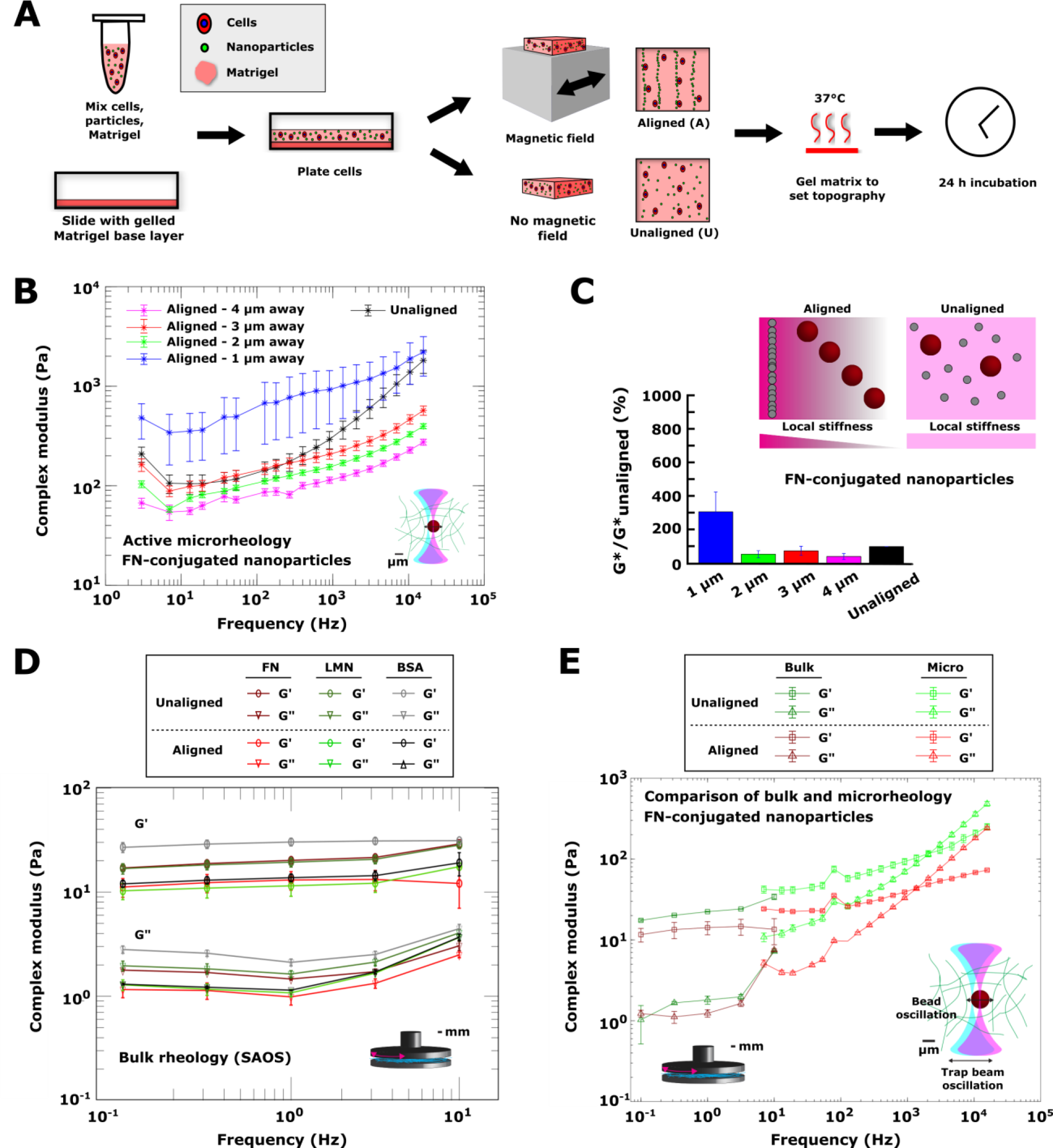
Characterization of the mechanical properties of engineered matrices across different length scales. (A) Schematic of 3D matrix patterning process. Human cells and superparamagnetic colloidal particles were suspended in Matrigel, plated on a glass slide containing a base layer of Matrigel, and either aligned in a magnetic field (aligned gels) or left unaligned and dispersed throughout the matrix (unaligned gels). Gels were then formed by heating at 37°C to set the matrix topography. Cells were fixed for analysis 24 h after seeding. (B) Complex modulus G* (mean ± SEM) vs. frequency curves obtained using optical trap-based microrheology in gels made with colloidal particles conjugated to human fibronectin. Moduli were measured at beads in unaligned gels (black), or in aligned gels at distances of 1 μm (blue), 2 μm (green), 3 μm (red), or 4 μm (pink) away from the nearest fiber. Samples were measured in triplicate, with at least 30 beads per sample analyzed. (C) Schematic and summary of microrheology experiments. In aligned gels, local stiffness increased closer to the fibers. In unaligned gels, stiffness was the same throughout the gel. Trend is evident by plotting the complex modulus (mean ± standard deviation), normalized to the complex modulus in unaligned gels, as a function of distance from the nearest fiber. The percentage was averaged across all measured frequencies for a given fiber distance to obtain the mean and standard deviation. (C) Bulk elastic (G’, circles) and viscous (G’’, triangles) components of complex moduli (mean ± SEM) of Matrigel gels made with colloidal particles coated in human fibronectin (red), tenascin C (green), or BSA (black), either unaligned (dark red, green, and black) or aligned (light red, green, and black) within a Matrigel matrix. Measurements were by parallel plate small angle oscillatory shear (SAOS) bulk rheology and were carried out in duplicate. (E) Elastic (G’,squares) and viscous (G’’, triangles) components (mean ± SEM) of complex moduli of gels made with colloidal particles conjugated to human fibronectin. Moduli were measured using either bulk rheology (dark green, dark red) or optical trap-based microrheology (light green, light red). For microrheology measurements, the moduli values at all distances from the nearest fiber were combined. Bulk rheology measurements were made in duplicate. For all microrheology measurements, samples were measured in triplicate, with at least 30 beads per sample analyzed.

**Figure 2.**
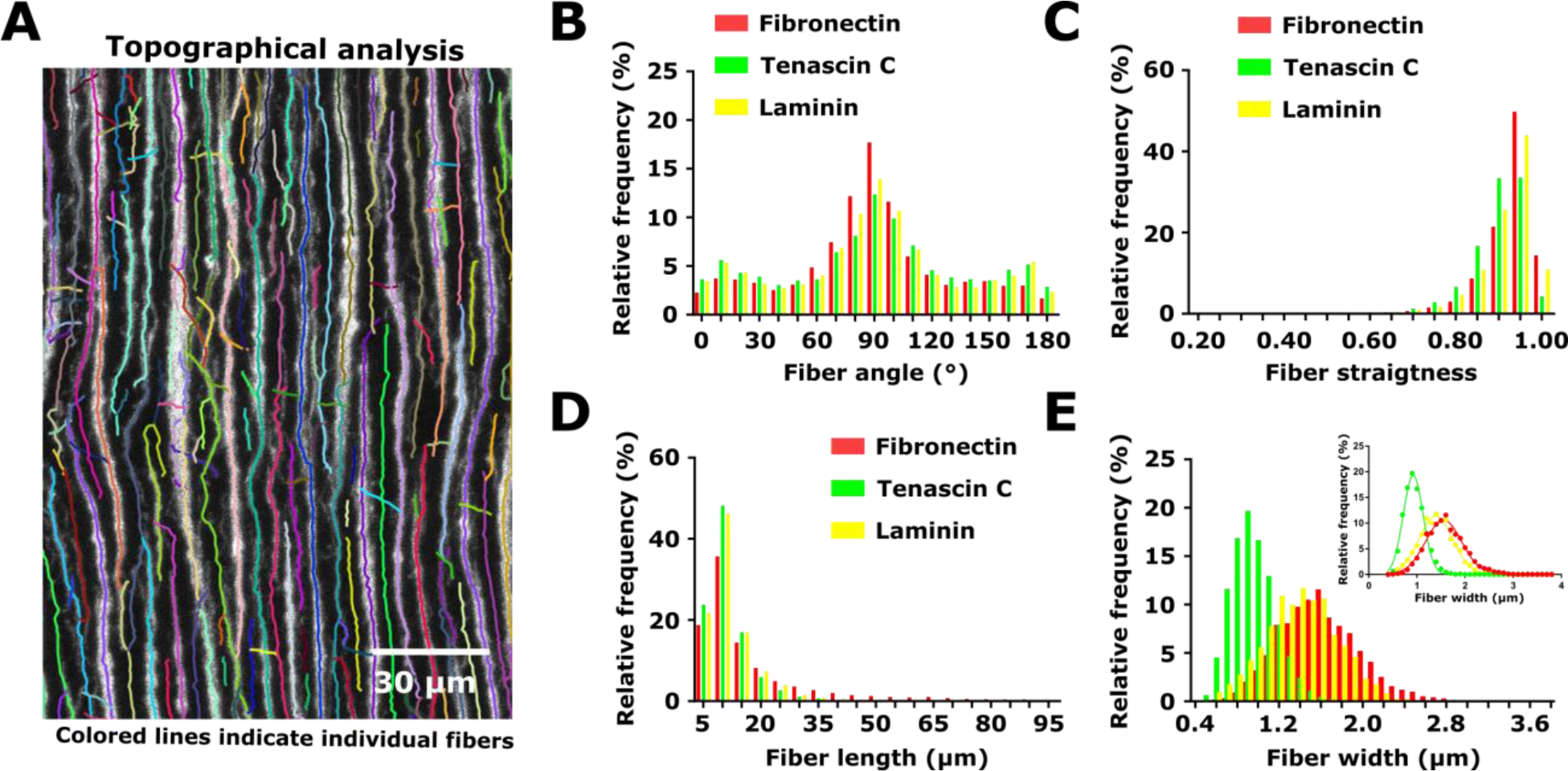
Characterization of the topographical properties of engineered matrices. (A) Representative image of fibers formed of fibronectin-conjugated colloidal particles and segmented using the ctFire fiber analysis toolbox. Colored lines indicate individual fibers segmented for analysis of fiber morphology. (B) Distribution of fiber angles of to the vertical in Matrigel matrices containing aligned fibers formed of fibronectin-, tenascin C-, or laminin-conjugated colloidal particles. (C) Distribution of fiber straightness values in Matrigel matrices containing aligned fibers formed of fibronectin-, tenascin C-, or laminin-conjugated colloidal particles. (D) Distribution of fiber lengths in Matrigel matrices containing aligned fibers formed of fibronectin-, tenascin C-, or laminin-conjugated colloidal particles. (E) Distribution of fiber widths in Matrigel matrices containing aligned fibers formed of fibronectin-, tenascin C-, or laminin-conjugated colloidal particles. Inset shows Gaussian fit of fiber width distributions. Three images from the same matrix were analyzed for aligned fibronectin-conjugated colloidal particles, whereas one image per matrix were analyzed for matrices containing aligned tenascin C- and laminin-conjugated colloidal particles. This resulted in analysis of 7288 fibers for fibronectin-conjugated particles, 4571 fibers for tenascin C-conjugated particles, and 3895 fibers for laminin-conjugated particles. Colored lines indicate individual fibers segmented for analysis of fiber morphology.

**Figure 3.**
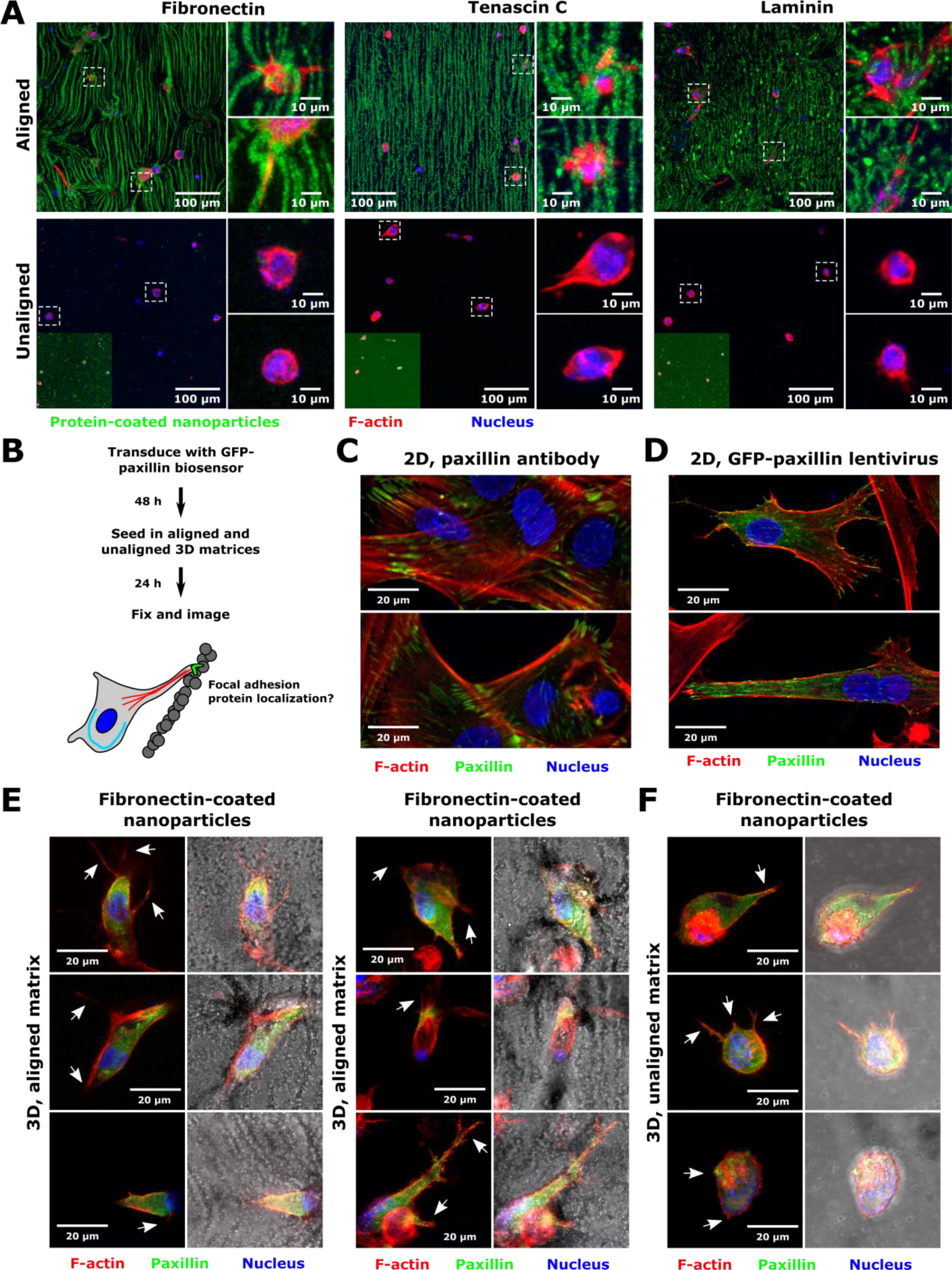
Human foreskin fibroblast (HFF) cell morphology in a 3D fibrillar matrix system. (A) Representative images of HFF cells embedded in aligned or unaligned Matrigel matrices containing colloidal particles conjugated to fibronectin, tenascin C, or laminin. In each panel, overview images are shown, with boxes to show detailed cell morphology (cell position in larger image indicated by dashed white boxes). In unaligned matrix images, insets show unaligned matrix with lookup table adjusted to show presence of dispersed fluorescent particles. Scales are indicated. (B) Schematic of experiment to assess focal adhesion protein localization in 3D matrices. Cells were transduced with a GFP-paxillin lentiviral biosensor prior to being embedded in aligned and unaligned Matrigel matrices. Cells were fixed, stained, and imaged after being embedded in matrices for 24 h. (C) Representative images of HFF cell focal adhesion formation after plating on two-dimensional tissue culture plastic, as assessed by immunofluorescent staining using a paxillin primary antibody. Images are single confocal slices. F-actin is displayed in red, paxillin in green, and the nucleus in blue. Scale is indicated. (D) Representative images of HFF cell focal adhesion formation after plating on two-dimensional surfaces coated with Matrigel, as assessed using the GFP-paxillin biosensor. Images are maximum intensity projections of confocal slices. F-actin is displayed in red, paxillin in green, and the nucleus in blue. Scale is indicated. (E) Representative images of HFF cells in aligned Matrigel matrices containing fibronectin-conjugated nanoparticles and expressing a GFP-paxillin biosensor. (F) Representative images of HFF cells in unaligned Matrigel matrices containing fibronectin-conjugated nanoparticles and expressing a GFP-paxillin biosensor. In panels (E,F), images are maximum intensity projections of confocal slices containing colloidal particles. F-actin is displayed in red, paxillin in green, and the nucleus in blue. Brightfield images are shown to illustrate particle alignment. Arrows indicate cell protrusions in the plane of the fibers. Scale is indicated.

### Protrusion analysis for cells embedded in 3D matrices

To analyze protrusion length and directionality, confocal tile scans acquired at 20x magnification and axial steps of 2 μm (1275.29 μm × 1275.29 μm, 1536 pixels × 1536 pixels) were opened in Fiji. Protrusions were measured only in the plane in which the fibers were in focus to quantify cell response to local fibers in aligned matrices, and only for cells clearly embedded in 3D for unaligned matrices. The line tool in Fiji was used to draw a line from the edge of the nucleus (manually identified) to the end of the protrusion to measure the protrusion length and protrusion angle. In aligned matrices, the line tool was also used to draw a line on the fiber immediately adjacent to the protrusion. The angle (0°-90°) between the protrusion and the neighboring fiber was calculated and recorded as the protrusion angle. For cells in unaligned matrices, the protrusion angle was calculated with respect to the vertical, and all angles were then mapped to be between 0° and 90°. For each cell type, matrix alignment status, drug, or antibody treatment, and nanoparticle ECM protein, 10-20 protrusions were measured from each 3D matrix. Statistical analysis and plot generation were done in Prism GraphPad 7. The number of matrices prepared for each condition is indicated in figure legends, and a matrix was used only if at least 10 cells were measured. Histograms were generated to visualize the distribution of protrusion angles in aligned and unaligned matrices. Protrusion lengths were compared between aligned and unaligned matrices using Sidak’s multiple comparisons test for a given ECM nanoparticle coating following two-way ANOVA. For mechanistic experiments, protrusion lengths were compared between control and treated cells for a given cell type by Dunn’s multiple comparisons post-test following a Kruskal-Wallis test.

### Analysis of matrix topography

Fibers were analyzed in Matrigel matrices containing colloidal particles conjugated to fibronectin, tenascin C, or laminin and HFF cells seeded for 24 h as described above. Analysis was performed on maximum intensity z projections of the fluorescent channel for the fibers in 425.10 μm × 425.10 μm (2048 pixels × 2048 pixels), 12-bit images acquired at 20× magnification as described above. Maximum intensity projections were performed on planes containing aligned fibers. Images were changed to 8-bit grayscale and analyzed using the ctFIRE V2.0 toolbox [30]. Minimum fiber length was set at 30 pixels, and max fiber width was set at 30 pixels. Three images from the same matrix were analyzed for aligned fibronectin-conjugated colloidal particles, whereas one image per matrix were analyzed for matrices containing aligned tenascin C- and laminin-conjugated colloidal particles. This resulted in segmentation and analysis of 7288 fibers for fibronectin-conjugated particles, 4571 fibers for tenascin C-conjugated particles, and 3895 fibers for laminin-conjugated particles.

### Fluorescence recovery after photobleaching

To generate matrices containing diffusible dextran, 430 μl of Matrigel was mixed with 12 μl of fibronectin- or BSA-conjugated colloidal particles (300 nm diameter) at 10 mg/ml and 50 μl of FITC dextran (fluorescein isothiocyanate dextran, average molecular weight 10 kDa, Sigma Millipore, Catalog #FD10S-250MG) at 984 μg/ml. This generated a mixture containing FITC dextran at 100 μg/ml. Proteins conjugated to colloidal particles were not fluorescently labeled so as not to interfere with the dextran fluorescence signal. To generate control Matrigel matrices lacking added colloidal particles, the same proportions were used, with 12 μl of serum free media replacing the 12 μl of particle solution.

Matrigel/particle/dextran mixtures were plated in wells of a 4-well chamber slide coated with 100 μl/well of Matrigel, as described above. Matrices were aligned on a magnet that was chilled in ice for 15 minutes as described above, or kept on ice away from the magnet for 15 minutes to generate unaligned matrices. The matrix was polymerized for 30 minutes at 37 °C. Matrices were then hydrated with 350 μl/well of serum free media containing 100 μg/ml of FITC dextran.

Fluorescence recovery after photobleaching (FRAP) experiments were carried out using the FRAP module on a Zeiss 780 confocal equipped with a Zeiss 20x Plan-Apochromat, 0.8 NA objective. One-photon, confocal, 12-bit, 2-dimensional images were acquired at lateral dimensions of 512×512 pixels to obtain a pixel size of 0.830 μm x 0.830 μm. A 50 pixel × 50 pixel square region of the matrices in the plane of the aligned fibers was bleached for ~30s and imaged every 500 msec for 240 cycles. Four scans of the region were acquired before bleaching. A reference region offset from the bleached region of the same dimensions was used to account for loss of signal due to repeated scanning.

FRAP parameters were calculated using the FRAP analysis module in Zeiss ZEN. Briefly, image intensity values over time were fit to the formula:

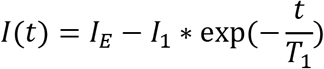

Curves were normalized to obtain mobile fractions and characteristic half maximum times using the built-in calculations in ZEN:

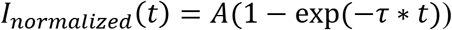

Where A = mobile fraction and the half maximum time is defined as:

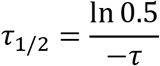

For each condition (particle protein coating and alignment status), three independent regions from two gels were measured. These measurements were grouped to obtain N=6 values prior to statistical comparisons. Analysis was performed in GraphPad Prism 7 using two-way ANOVA comparing aligned and unaligned matrices containing either fibronectin- or BSA-conjugated colloidal particles, followed by Sidak’s multiple comparisons test between conditions for a given ECM protein.

### Bulk rheology

Small angle oscillatory shear bulk rheology measurements were carried out at the Georgetown University Institute for Soft Matter Synthesis and Metrology using an Anton Paar Physica MCR 301 rheometer equipped with a PP-25 measuring plate (parallel, 25 mm diameter). Gels were polymerized on 50 mm glass bottom dishes (Wilco, Amsterdam, The Netherlands, Catalog #GWSB-5040). Samples were prepared with 120 μl Matrigel on bottom of Wilco dish polymerized prior to the Matrigel/bead matrix. To this, a mixture of 430 μl Matrigel, 30 μl SFM, 12 μl paramagnetic beads, and 20 μl of 1×10^5^/ml polystyrene beads (2×10^6^ beads total). Beads were 1 μm rhodamine carboxylated fluorospheres (ThermoFisher, Catalog #F8821).

Samples were either aligned or unaligned as described above and were hydrated with a superlayer of media for storage and transport. Media was removed with a pipette before measurements. The instrument achieved contact with the sample with a trigger force of 0.1 N normal and the excess gel was trimmed around the plate to ensure proper contact boundary conditions. The complex modulus was measured at 1% strain at frequencies 0.1-10 Hz. Measurements were carried out in duplicate.

### Optical tweezer-based microrheology

Samples were prepared in Wilco dishes identically to those made for bulk rheology measurements prior to characterization via optical tweezer-based microrheology. For complete experimental details, see [31–33]. Our home-built setup consists of a 1064 nm trapping beam steered by an acousto-optic deflector to oscillate the trap and a stationary 975 nm detection beam that is coupled into and colocated with the trap with a dichroic before being sent into the backport of an inverted microscope with a long working distance water objective and a high NA condenser. Telescope lenses conjugate the optical plane at the acousto-optic deflector (AOD) to the back aperture of the condenser, which is placed in Kohler illumination after the object is focused in the specimen place. Above the condenser, the detection beam is relayed to a quadrant photodiode for back focal plane interferometric position detection. Each bead is positioned precisely in the center of the trap by scanning it through the detection beam in three dimensions using a piezo nanopositioning stage while recording the voltages from the QPD. The V-nm relation of the QPD is calibrated in situ by fitting the central linear region of the detector response to scanning the bead through the detection beam in the direction of the oscillations, giving β in V/nm (stuck bead method). A second QPD records the position of the trapping laser to find the relative phase lag between the bead and trap oscillations. The optical trap stiffness *k* is determined in situ from the thermal power spectrum of each measured bead while the trap is stationary, using the active-passive calibration method[34]. Together with *β, k*, and the bead’s mass *m* and radius *a*, the trajectories yield the complex modulus as a function of frequency, G*(ω), of each bead’s surrounding microenvironment. In this equation, the complex modulus, G*(ω), can be broken down into components, with *G* *(*ω*) = *G*’(*ω*) + *iG*”(*ω*), where the real part, G’(ω), is the elastic component and the imaginary part, G’’(ω), is the viscous component. The complex modulus, G*(ω), is calculated as 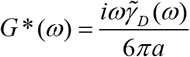, where the friction relaxation spectrum 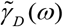 is related by the equation 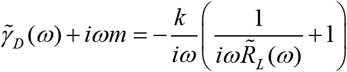 to the active power spectrum 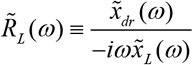, with 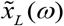 and 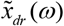 the Fourier transforms of the time series of the positions of the trapping laser and the driven bead respectively, recorded while the trap is oscillating. The stifness 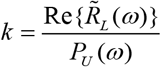 is determined from the real part of the active power spectrum and the passive power spectrum, 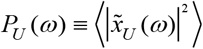, where 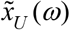 is the Fourier transform of the time series of the undriven bead’s thermally fluctuating position while the trap is held stationary. Each bead is subjected to fourteen consecutive 2 s pulses, with the trap alternately oscillating or stationary. Amplitude of oscillations was set to 20 nm with power of 100 mW at the back aperture. Only probes at distances exceeding ~30 μm away from the cover slip surface to minimize drag in consideration of Faxen’s law were measured [35].

Samples were measured in triplicate with at least 30 beads per sample measured. Laser power was set to 100 mW at the back aperture. Data were analyzed using custom MATLAB programs. Experiments were controlled using custom LabVIEW programs. In Figure 1C, for each distance from the nearest fiber and each measured frequency, the mean complex modulus was divided by the mean complex modulus in unaligned gels, and moduli are expressed as percentages of the unaligned gel complex modulus. These percentages were averaged across all frequencies to obtain the plotted means and standard deviations.

### Cell seeding on 2D silicone substrates of known elastic modulus

Cytosoft 6-well plates (Advanced BioMatrix, San Diego, CA, Catalog #5190-7EA; 0.5 kPa, 2 kPa, and 64 kPa elastic modulus) or standard 6-well tissue culture plastic dishes were incubated with human plasma fibronectin (Millipore Sigma, Catalog #FC010) at a concentration of 10 μg/ml diluted in PBS (3 ml/well of ECM protein solution added) for 1 h at room temperature. The coating solution was aspirated, and wells were washed twice with PBS. HFF or U87 cells were washed with phosphate buffered saline (PBS), detached from the cell culture flask using 10 mM EDTA in PBS, washed once in growth medium, centrifuged at 1000 rpm for 5 min, and resuspended to a concentration 100,000 cells/ml in serum free medium. To each well, 1 ml of the cell suspension (100,000 cells total) and 1 ml of serum free media were added. Plates were incubated overnight at 37°C prior to fixation and staining.

For samples plated on substrates of varying stiffness, culture media was aspirated, and 3 ml of 4% paraformaldehyde diluted in PBS was added to each well. Samples were fixed at room temperature for 15 min and washed twice with PBS. Cells were then permeabilized with 0.1% Triton X-100 in PBS for 5 min and washed twice more with PBS. A phalloidin stock solution was prepared by dissolving 10 nmol of phalloidin-Atto 488 (Millipore Sigma, Catalog #49409-10NMOL) in 500 μl methanol. To fluorescently label F-actin and the nucleus, a solution containing 1 μg/ml Hoechst 33258 and 20 μl phalloidin stock/ml in 1% BSA in PBS was prepared. Cells were stained with 500 μl staining solution/well for 1 h at room temperature. Finally, wells were washed twice with PBS and stored in PBS prior to imaging. For each condition, two wells per cell type were prepared simultaneously.

For cells plated on 2D silicone substrates, imaging was carried out at 20× magnification using a ThermoFisher EVOS FL Cell Imaging System. Four random fields of view were selected for each well. Two wells were prepared and imaged simultaneously for each cell type and substrate stiffness. A minimum of 73 cells were imaged and analyzed for each condition.

### Analysis of cell morphology on 2D substrates

Cell morphology on 2D substrates of varying elastic modulus was analyzed using Fiji. Images obtained from the F-actin channel were binarized using the default Fiji settings and processed using the “Close-” function in Fiji. Dead and truncated cells were removed from the images, and shapes were manually separated or joined when necessary to more closely match the original images. Another investigator compared the final drawings to the original images for independent verification of the results. Circularity and area were calculated from the binarized images using the “Analyze Particle” function in Fiji. Measurements were made on cells in four random fields of view for each well, with two wells per substrate elastic modulus prepared and imaged simultaneously for each cell type. Statistics were carried out and plots were generated in GraphPad Prism 7. The effects of cell type and matrix stiffness on cell area and aspect ratio were analyzed using two-way ANOVA with Tukey’s multiple comparisons post-tests between all combination of substrate stiffnesses for a given cell type.

### Zebrafish experiments

Animal studies were conducted under protocols approved by the National Cancer Institute and the National Institutes of Health Animal Care and Use Committee. Transgenic Tg(fli:EGFP) zebrafish [36] were maintained on a 14-hour light/10-hour dark cycle according to standard procedures at 28.5°C. Larvae were obtained from natural spawning, raised at 28.5°C, and maintained in fish water (60 mg Instant Ocean^©^ sea salt [Instant Ocean, Blacksburg, VA] per liter of DI water). Larvae were checked regularly for normal development. For all experiments, larvae were transferred to fish water supplemented with N-phenylthiourea (PTU; Millipore Sigma, Catalog #P7629-25G) between 18-22 hours post-fertilization to inhibit melanin formation. PTU water was prepared by dissolving 16 μl of PTU stock (7.5% w/v in DMSO) per 40 ml of fish water. Water was replaced daily.

For cell injections, HFF and U87 cells were detached from cell culture plates using 10 mM EDTA, stained with CellTracker Deep Red membrane dye (ThermoFisher, Catalog #C34565) diluted to 1 μM in PBS for 30 min at 37°C, washed with PBS, and resuspended to a concentration of 1 million cells/20 μl PBS. An anesthetic of buffered tricaine was prepared by adding 4.2 ml tricaine stock (400 mg tricaine powder [Millipore Sigma, Catalog #E10521-50G], 97.9 ml deionized water, 2.1 ml of 1 M Tris) per 100 ml of fish water supplemented with PTU. 2 days post-fertilization (dpf) Tg(fli:EGFP) larvae were anesthetized and oriented on an agarose bed. A volume of 2-5 nl of the cell suspension (~100-250 cells) were injected to the zebrafish hindbrain. Zebrafish were maintained for 24 h following injection at 33°C in fish water supplemented with PTU or water containing 100 μM blebbistatin or vehicle control (DMSO).

For imaging, zebrafish were anesthetized and mounted in a lateral orientation in 1% agarose in a Lak-Tek 4-well chambered coverglass well. Tricaine water was added to the well prior to imaging. Images were obtained on a Zeiss 780 LSM confocal microscope. One-photon, confocal, 12-bit, 2-dimensional images were acquired at lateral dimensions of 512×512 pixels with a Zeiss 20× Plan-Apochromat, 0.8 NA objective, and images at axial distances of 1 μm were stacked to obtain three-dimensional datasets. Images presented in Figure 6 are average intensity projections.

## RESULTS

### Characterization of the bulk topographical and rheological properties of engineered 3D aligned matrices

We used our recently developed platform in which we incorporate a number of human proteins into the three-dimensional fibrillar architecture system [17, 37]. ECM proteins were labeled with a fluorescent marker and conjugated to 300 nm-diameter carboxylated superparamagnetic particles. To generate 3D matrices with embedded cells and fibers, cells were mixed with Matrigel and colloidal particles and plated in coverslip-bottom chamber slides that had been previously covered with a thin layer of gelled Matrigel. The slide was then either placed on a magnet to generate aligned fibrils or kept far from the magnet to maintain a random dispersion of particles. Finally, the slide was kept at 37°C to gel and set the nanoparticle topography, and cells were incubated in the presence of serum free medium for 24 h (**Figure 1A, Supplementary Figure 1A-C**). Aligned and unaligned matrices contained the same absolute amount of ECM proteins conjugated to colloidal particles and the same number of colloidal particles per unit volume within the hydrogel, differing only in the alignment status of the colloidal particles.

Cellular migration and cell morphology are sensitive to the compliance and viscoelasticity of their immediate milieu [2, 19]. We thus characterized the mechanical properties of both aligned and unaligned laminin-rich ECM (Matrigel) matrices in the absence of cells. Macroscale mechanics are often determined from models where the material can be assumed to act as a continuum. In order for the continuum assumption to apply, the characteristic length scale of underlying structural components must be much smaller than that of the physical measurement [38]. One concern relevant to characterization of the composite matrices is that the magnetic particles forming the aligned fibrils are rigid, and that this property could in turn affect cell phenotype. Thus, we reasoned that at the microscale, the self-assembled fibrils may give rise to local variations in mechanics that will not be resolved with bulk rheology. We performed optical tweezer-based active microrheology to probe heterogeneous mechanical properties in 3D microenvironments on the length scales of cellular mechanotransduction (~micron) (**Figure 1B,C**). For these experiments, we focused on composite Matrigel gels containing fibronectin-conjugated colloid particles. Active microrheology measurements revealed a gradient in the complex modulus (G*) as a function of distance from the assembled fiber (**Figure 1B,C; Supplementary Figure 2A**). Regions within 1 μm of the fibers, which were comprised of rigid paramagnetic particles, were stiffer than the unaligned gels, with G* over 3-15,000 Hz ranging from 0.5 − 2.5 kPa and 0.2 − 2.0 kPa for aligned and unaligned gels, respectively, with both frequency (p = 2.8e−80) and fiber alignment (p = 1.5e−39) having statistically significant effects on G* by two-way ANOVA (**Figure 1B,C)**. At further distances of 2–4 μm from the nearest fiber, local stiffness decreased (**Figure 1B,C**). Complex modulus values at distances of 2–4 μm from the nearest fiber were slightly less those than in unaligned gels (**Figure 1B,C**). The complex modulus at a frequency of 3 Hz was 65 Pa, 100 Pa, and 175 Pa for beads at distances 2 μm, 3 μm, and 4 μm, respectively, while the complex modulus at 15,000 Hz was 300 Pa, 400 Pa, and 600 Pa for these distances. The effect of distance from the fiber on complex modulus across frequencies is summarized in **Figure 1C**.

We determined that there is a difference in frequency dependence of the complex moduli in aligned vs. unaligned gels (**Figure 1B**; **Supplementary Figure 2**). To quantitatively assess the frequency dependence of the complex moduli in these gels, we fit the complex modulus using a power law model, G*(ω)=Aω^b^, where ω is the frequency. For beads 1–4 μm away from an aligned fiber, the frequency dependence was similar, with G* weakly dependent on frequency, having power law fits of – 1 μm: A=194 (95% C.I.: 147, 241), b=0.24 (95% C.I.: 0.21, 0.27), r^2^: 0.95; 2 μm: A=35 (95% C.I.: 24, 36), b=0.24 (95% C.I.: 0.20, 0.28), r^2^: 0.92; 3 μm: A=42 (95% C.I.: 26, 59), b=0.25 (95% C.I.: 0.21, 0.30), r^2^: 0.90; 4 μm: A=30 (95% C.I.: 21, 38), b=0.22 (95% C.I.: 0.18, 0.25), r^2^=0.91). In unaligned gels, the power law exponents were closer to the value (0.75) predicted for semi-flexible polymers (A=5 (95% C.I.: 2, 8), b=0.61 (95% C.I.: 0.54, 0.68), r^2^: 0.98). In addition, the frequency dependence of the elastic and viscous contributions to the complex moduli (G* = G’ + *i*G”, where G’ = elastic component and G” = viscous component) differed between the aligned and unaligned gels, with crossover frequencies (at which G’’ first exceeds G’) of ~1 kHz in unaligned gels and ~10 kHz in aligned gels (**Supplementary Figure 2**). As an example, the crossover frequency in aligned matrices at a distance of 3 μm from the nearest fiber was much greater than that in unaligned gels (**Supplementary Figure 2B**).

To assess how micron-scale rheological measurements compared to bulk rheology, we characterized bulk mechanical properties of aligned and unaligned Matrigel hydrogels using parallel plate small angle oscillatory shear (SAOS) bulk rheology (**Figure 1D**). Bulk rheological measurements revealed that the complex moduli (G* = G’ + *i*G”, where G’ = elastic component and G” = viscous component) of aligned vs unaligned (random) matrices were comparable independently of the ECM coating used **(Figure 1D).** These hydrogels were mostly elastic, where the shear elastic moduli ranged from 10-30 Pa with very little viscous component over the range of ~0.1-100Hz (**Figure 1D**), less rigid than the ~100 Pa stiffness reported for collagen gels [39]. The complex moduli of Matrigel matrices containing colloidal particles were similar to the bulk rheological properties of Matrigel in the absence of colloidal particles that we have reported previously [17, 31]. In the low frequency regime, bulk and microrheology methods gave nearly identical values for the complex modulus (**Figure 1E**). Fluorescence recovery after photobleaching (FRAP) measurements of aligned and unaligned Matrigel matrices containing colloidal particles conjugated to fibronectin or BSA revealed a slight decrease (~15%) in both the mobile fraction and half-max time in aligned vs. unaligned gels (**Supplementary Figure 3**). Values in unaligned matrices were nearly identical to those obtained in Matrigel lacking colloidal particles. While alignment status was a significant sources of variation in FRAP parameters by two-way ANOVA, half-maximum times in both aligned and unaligned gels were ~7-8 seconds, much shorter than the overnight cell spreading times in protrusion experiments (**Supplementary Figure 3**).

We next characterized the topographical properties of the engineered matrices. Matrigel matrices were seeded with human foreskin fibroblasts (HFFs) and colloidal particles conjugated to human fibronectin, tenascin C, or laminin, aligned, incubated for 24 h, and imaged. Individual fibers were analyzed using the ctFire analysis toolbox [30] (**Figure 2A, Supplementary Figure 4**). Fibers were oriented primarily in one direction by the magnetic field (**Figure 2B**) and exhibited low curvature, regardless of the ECM protein used to coat the colloidal particles (**Figure 2C**). Fiber lengths were slightly longer for fibronectin- vs. tenascin C- and laminin-coated colloidal particles, but for all ECM proteins, the length of an individual fiber was typically ~10 μm (**Figure 2D**). Fibers were typically ~1-2 μm in width, though fibers formed with tenascin C-coated colloidal particles were somewhat thinner than those formed from colloidal particles formed with fibronectin- or laminin-coated particles, as observed in Gaussian fits of the fiber width distributions (**Figure 2E**).

### Human foreskin fibroblasts and glioblastoma cells respond to aligned fibrils for a myriad of human ECM proteins in the 3D microenvironment

Having characterized the physical properties of the engineered matrices, we examined cellular response to aligned topographical cues using human HFF and U87 cells in Matrigel matrices. Protrusions serve as sensors of the local environment. Previous work has demonstrated that cells protrude along aligned topographical cues in collagen gels [40] and on microcontact-printed surfaces [41], and we therefore assessed whether similar contact guidance was observed in our engineered matrices. Protrusions in response to fibrillar topography were quantified in the presence of colloidal particles conjugated to human fibronectin, tenascin C, and laminin prior to being embedded within Matrigel. In aligned matrices, both cell types were spindle shaped and formed long, actin-rich protrusions, as indicated by staining with phalloidin (displayed in red; **Figure 3A**). In these aligned matrices, large (on the order of mm) areas could be patterned to contain oriented fibers with exposed ECM proteins for cell binding (**Supplementary Figure 5**). Cells embedded in matrices containing unaligned colloidal particles (and thus, the same absolute amount of human ECM proteins) remained largely spherical and had shapes similar to that of cells in Matrigel alone (**Figure 3A; Supplementary Figure 6A,B**). Additionally, the engineered alignment method was amenable to the production of alignment in other matrices, including agarose and hyaluronic acid (**Supplementary Figure 7**). Some particle aggregation was observed in unaligned matrices, but in general, the fluorophore density of the dispersed colloidal particles was not sufficient to generate a fluorescent signal. In **Figure 3A** and **Supplementary Figures 6** **and 7**, insets in images of unaligned matrices show these matrices with the lookup tables adjusted to demonstrate the presence of dispersed nanoparticles in the hydrogels.

We next asked if mature focal adhesions are formed in the aligned 3D Matrigel matrices (**Figure 3B**). We transduced HFF cells with GFP-paxillin. As a control, we assessed focal adhesion formation first on 2D surfaces, where immunofluorescence was performed with an antibody directed against paxillin. Fluorescence images revealed large focal adhesions at the distal ends of cell protrusions (**Figure 3C**). We confirmed similar distributions for cells transduced with the paxillin biosensor, where images revealed both large adhesions and cytoplasmic GFP-paxillin (**Figure 3D**). When transduced cells were introduced to Matrigel matrices containing aligned fibrils formed of fibronectin-conjugated colloidal particles, paxillin expression was diffuse, and we noted numerous instances where protrusions parallel or perpendicular the fibrils lacked plaques of paxillin (**Figure 3E**), while some protrusions had small paxillin-containing adhesions (**Figure 3E**, bottom right panel). Paxillin expression was similarly diffuse in unaligned 3D matrices (**Figure 3F**).

To quantitate the response of normal and malignant human cells in the engineered fibril system, HFF and U87 cells were fixed and imaged after being embedded in aligned or unaligned matrices (or Matrigel control without additional particles), and protrusion length and angle compared to local fiber alignment were quantified (**Figure 4A,D**). HFF cells plated in aligned matrices preferentially sent out protrusions along or perpendicular to the fibers (**Figure 4A**). In contrast, cells remained spherical in unaligned matrices and extended thin, randomly oriented filopodia-like protrusions (**Figure 4A,** right panel). Time-lapse videos of cells embedded in aligned matrices revealed that cells typically sent out protrusions along or perpendicular to fibers before locally contracting the matrix (**Supplementary Videos 1,2**). The length of protrusions of cells in aligned matrices 24 h after seeding significantly increased compared to cells in unaligned matrices, where protrusion lengths were similar to those seen in Matrigel alone (**Figure 4B**). The local angle between fibers and cell protrusions in aligned matrices was preferentially either 0° or 90° (**Figure 4C,** left panel), indicating protrusions sent out parallel or perpendicular to the fibers (see insets in **Figure 4A**). In unaligned matrices, protrusions were randomly distributed around the cell body (**Figure 4C,** right panel). These trends held across all of the ECM protein coatings tested, suggesting that the presence of a topographical cue was more important in the observed cell extensions than specific integrin-ECM interactions for fibers formed in a Matrigel base matrix. Although both ECM coating and matrix alignment were significant sources of variation in protrusion length by two-way ANOVA, matrix alignment status was the predominant factor, accounting for 46% of the observed variation, and protrusions were significantly longer in aligned vs. unaligned matrices for all ECM proteins tested. The assembly of ECM protein-conjugated colloidal particles into fibers could increase the local effective ligand density in the vicinity of the cell compared to unaligned gels. Thus, we also conjugated bovine serum albumin (BSA), a protein lacking binding moieties for cell adhesion proteins, to the colloidal particles and formed aligned and unaligned gels (**Supplementary Figure 6C**). Cell response (increased protrusion length and protrusion alignment to local fibers) was identical in this case (**Figure 4B,C)**. We also observed that paxillin expression was diffuse for cells cultured in matrices containing BSA-conjugated particles, similar to what was observed in aligned matrices containing fibronectin-conjugated particles (**Supplementary Figure 6D**).

**Figure 4.**
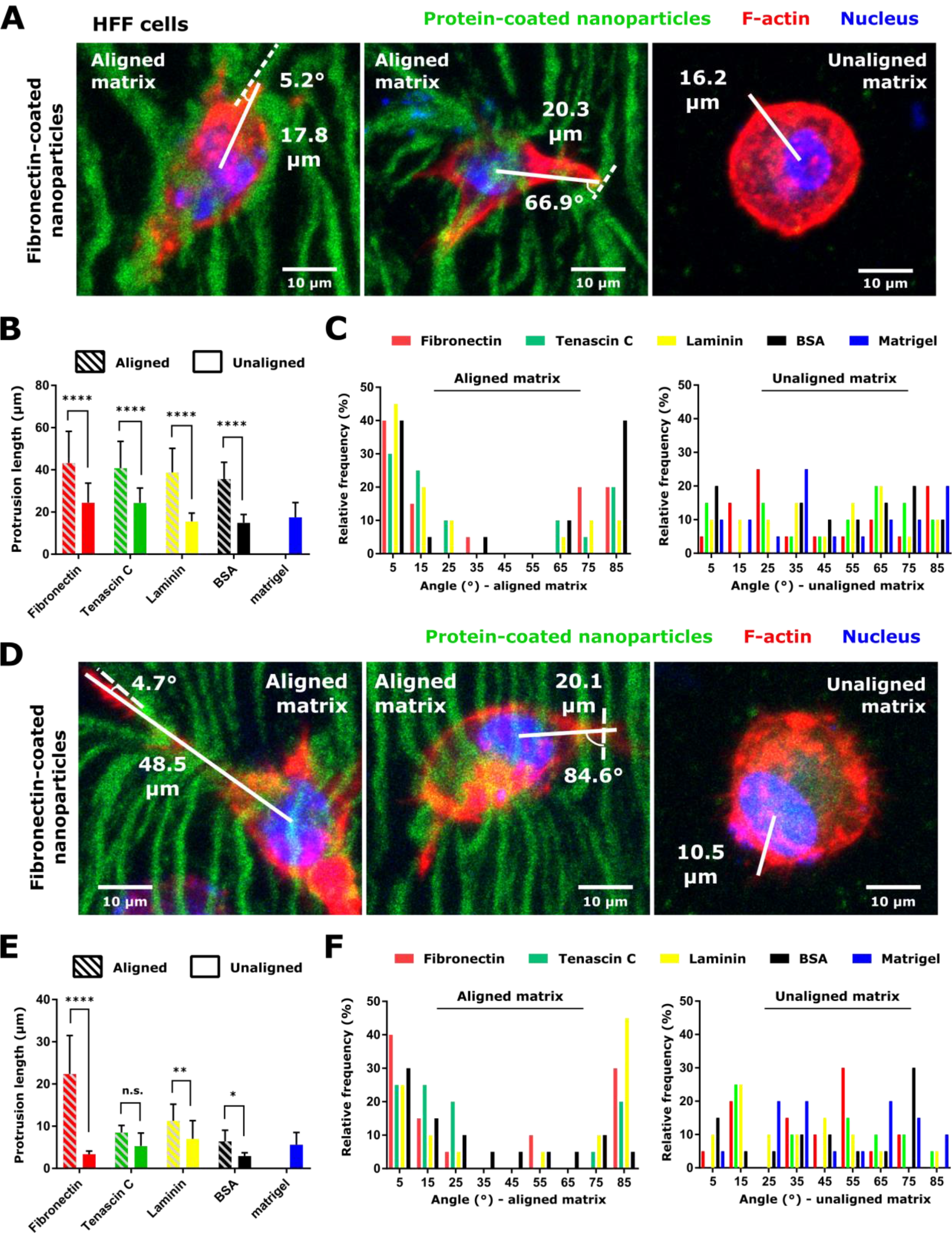
Increased cell protrusion generation and preferential protrusion orientation in aligned vs. unaligned matrices for HFF and U87 cells. (A) Representative images of HFF cells embedded in an aligned and unaligned Matrigel matrix containing colloidal particles conjugated to fibronectin. Images show detailed morphology of cells with measured protrusion lengths and protrusion angles. (B) Protrusion length (mean ± standard deviation) of HFF cells as a function of matrix alignment status and colloidal particle ECM protein conjugation. Average protrusion length in Matrigel lacking colloidal particles is also shown. ****, p<0.0001 by Sidak’s multiple comparisons test following two-way ANOVA. (C) Relative frequency distribution of the angle between HFF protrusions and nearest neighbor fibers (left panel) and in unaligned matrices (right panel) for Matrigel matrices containing colloidal particles conjugated to human fibronectin, tenascin C, laminin, or BSA. Protrusion direction distribution in Matrigel lacking colloidal particles is also shown. For each cell type, matrix alignment status, and nanoparticle ECM protein, 20 protrusions were measured from a single 3D matrix. (D) Representative images of U87 cells embedded in an aligned and unaligned Matrigel matrix containing colloidal particles conjugated to fibronectin. Images show detailed morphology of cells with measured protrusion lengths and protrusion angles. (E) Protrusion length (mean ± standard deviation) of U87 cells as a function of matrix alignment status and nanoparticle ECM protein conjugation. Average protrusion length in Matrigel lacking colloidal particles is also shown. *, p<0.05; **, p<0.01; and ****, p<0.0001 by Sidak’s multiple comparisons test following two-way ANOVA. (F) Relative frequency distribution of the angle between U87 protrusions and nearest neighbor fibers (left panel) and in unaligned Matrigel matrices (right panel) for matrices containing colloidal particles conjugated to human fibronectin, tenascin C, laminin, or BSA. Protrusion direction distribution in Matrigel lacking colloidal particles is also shown. For each cell type, matrix alignment status, and nanoparticle ECM protein, 20 protrusions were measured from a single 3D matrix. In panels (A,D), colloidal particles are displayed in green, F-actin in red, and the nucleus in blue. Images are maximum intensity projections of confocal slices containing aligned fibers, or of cells embedded in 3D. Scales are indicated.

We next asked if U87 cells showed similar behavior to the normal cell line. Similar responses to matrix alignment for the diversity of ECM chemistries tested where U87 cells generated protrusions preferentially parallel or perpendicular to local fibers and remaining largely spherical in unaligned matrices (**Figure 4D**). U87 cells extended longer protrusions in aligned than unaligned matrices for a given ECM protein (**Figure 4E**). Similar to the case seen in HFF cells, both ECM protein and alignment status were significant sources of variation in protrusion length by two-way ANOVA, but alignment status again accounted for the largest percentage of the variation observed. Additionally, topographical cues in aligned matrices led to U87 cell protrusions predominantly parallel or perpendicular to local fibers (**Figure 4F,** left panel). These trends were not observed in unaligned matrices, where protrusions were again distributed around the cell body instead of in preferred directions (**Figure 4F,** right panel).

Having determined that there were local heterogeneities in microscale mechanics in aligned Matrigel matrices and that cells generated directional protrusions in response to alignment, we aimed to assess how environmental stiffness regulated cell elongation. We thus characterized cell phenotypic response to substrates of differing mechanical properties. Previous work has shown that sensitivity to ECM substrate stiffness is dependent on the concentration of ECM ligand [16, 42], and we therefore compared cell morphology on substrates coated with a comparable concentration of ECM protein as that conjugated to the colloidal particles (approximately 0.3 μg/cm^2^) with stiffnesses that spanned several orders of magnitude from 0.5 kPa - >GPa (**Supplementary Figure 8A**). This dynamic range allowed us to probe responses across the stiffnesses of several tissues, from the brain to the bone. We determined that the cell morphology and the presence of actin cytoskeletal structures in HFF and U87 cells were largely insensitive to the stiffness of the underlying ECM substrate until stiffnesses reached the GPa range, well higher than the difference in stiffness observed near and far from aligned fibers in the engineered system. Substrate stiffness was not a significant source of variation in cell aspect ratio by two-way ANOVA (**Supplementary Figure 8B**), and significant differences in areas for U87 cells were only observed between the much stiffer tissue culture plastic compared to the softer silicone substrates (**Supplementary Figure 8C**). HFF cell spreading was also reduced at the softest substrate stiffness (0.5 kPa) compared to intermediate stiffness values (**Supplementary Figure 8C**).

### Integrin β1 and fascin contribute to cell protrusion generation in aligned matrices

Having observed cell protrusions similar to elongated filopodia in response to topographical cues in the presence of a local gradient in micromechanics, we set out to understand mechanistic drivers of cell response within 3D substrates by interrogating processes implicated in protrusion generation (**Figure 5A,B**). We focused on Matrigel matrices containing aligned fibers formed of fibronectin-conjugated colloidal particles. Cells use integrin-rich filopodia adhesions to sense ECM gradients. We thus reasoned that integrin β1 may be a key regulator needed for the observed elongation and alignment of cellular protrusions. Silencing integrin β1 using siRNA significantly reduced the average length of protrusions formed for both HFF and U87 cells in aligned 3D matrices (**Figure 5B-C, Supplementary Figure 9A**). However, the protrusion angle relative to the angle of the nearest fiber was not changed upon knockdown (**Figure 5D,E**). Similar results were obtained upon treatment with a function-blocking antibody directed against integrin β1 (**Supplementary Figure 9B-D**). Extension of a filopodium is driven by a fascin-dependent bundling of actin filaments. Inhibition of fascin, using the inhibitor fascin-G2 (**Figure 5A,B**), had a similar effect as knockdown of integrin β1, shortening protrusion lengths while not affecting protrusion angles (**Figure 5F-H**). In summary, inhibition of integrin β1 and fascin tended to result in cells remaining fairly rounded in aligned matrices (**Figure 5B**).

**Figure 5.**
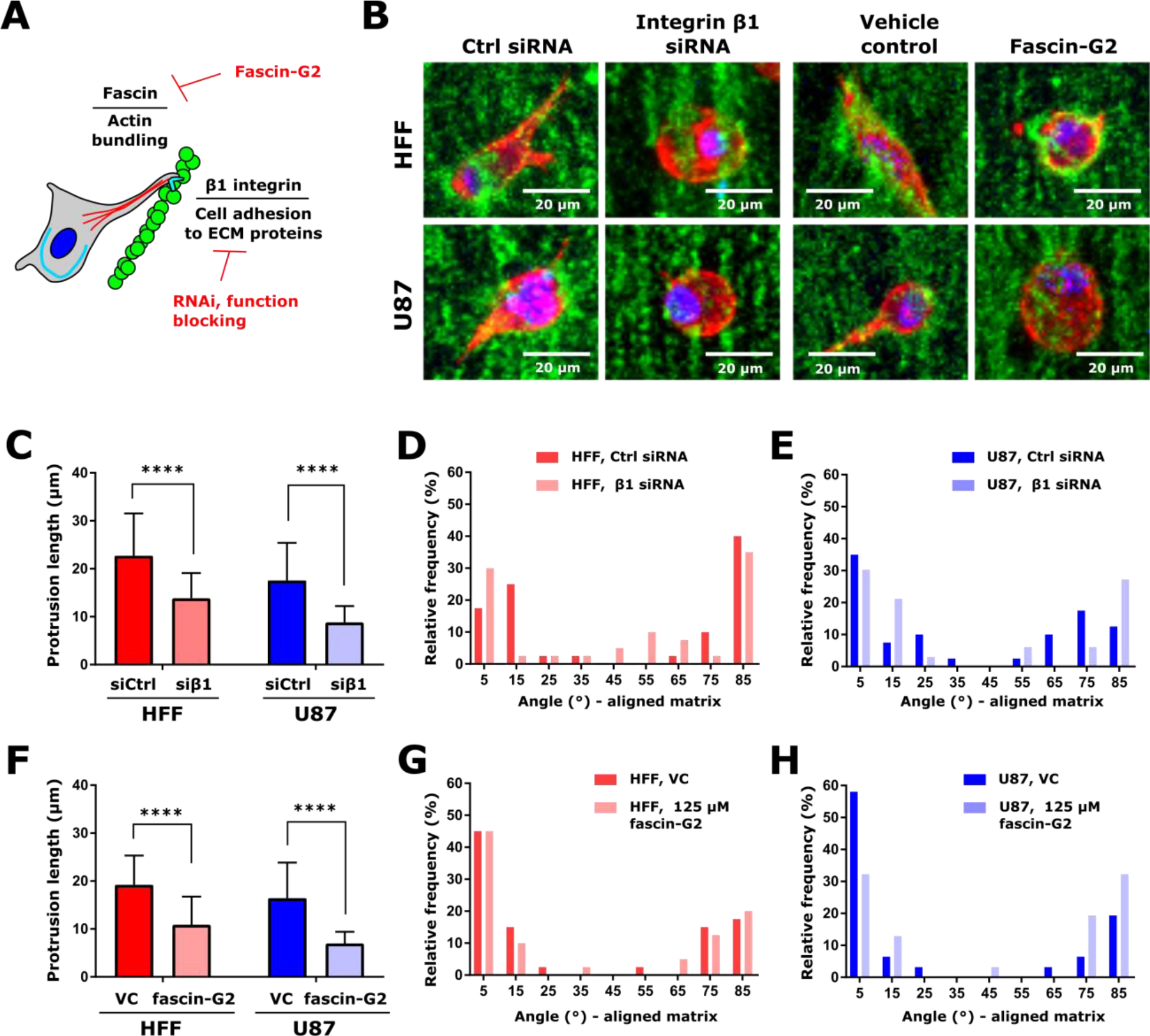
Modulation of protrusion generation via inhibition integrin β1 activity and actin bundling. (A) Schematic of cellular processes tested for effect on protrusion generation. Integrin adhesions were mediated via knockdown of integrin β1 and application of a function-blocking antibody, and actin bundling via fascin was inhibited by treatment with the fascin inhibitor fascin-G2. (B) Representative images of HFF and U87 cells in Matrigel matrices containing aligned fibronectin-containing particles upon knockdown of integrin β1 and application of 125 μM fascin-G2, or the appropriate controls. Particles are displayed in green, F-actin is displayed in red, and the nucleus is displayed in blue. Scale is indicated. (C) Cell protrusion length (mean ± standard deviation) in cells transfected with siRNA targeting integrin β1 (siβ1) or non-targeting control siRNA (siCtrl). (D) Distribution of angles between cell protrusions and the nearest fiber for HFF cells transfected with siRNA targeting integrin β1or non-targeting control siRNA. (E) Distribution of angles between cell protrusions and the nearest fiber for U87 cells transfected with siRNA targeting integrin β1or non-targeting control siRNA. In panels C-E, two matrices were analyzed per cell type and treatment. This resulted in analysis of 40 protrusions per condition for HFF cells and 40 (Ctrl siRNA) or 33 (integrin β1 siRNA) protrusions analyzed for U87 cells. (F) Cell protrusion length (mean ± standard deviation) in cells treated with 125 μM fascin-G2 or VC. (J) Distribution of angles between cell protrusions and the nearest fiber for HFF cells treated with 125 μM fascin-G2 or VC. (G) Distribution of angles between cell protrusions and the nearest fiber for U87 cells treated with 125 μM fascin-G2 or VC. In panels (F-H), 20 protrusions were analyzed from two matrices per cell type and treatment, resulting in 40 protrusions per condition measured. In panels (C,F), protrusion length was analyzed by Kruskal-Wallis test with Dunn’s multiple comparisons test within each cell type. ****, p<0.0001.

### Myosin II contributes to cell alignment in response to local topographical and stiffness gradients both in vitro and in vivo

As myosin II has been implicated in cell mechanosensing [25], we treated cells with the myosin II inhibitor, blebbistatin, and quantified cellular protrusions in aligned Matrigel matrices containing fibronectin-conjugated particles (**Figure 6A-E**). Upon treatment with blebbistatin, cells embedded in the matrices formed numerous long, spindly protrusions that were not well-aligned to the fibers (**Figure 6B**), and average protrusion length upon blebbistatin treatment remained unchanged in HFF cells and increased in U87 cells (**Figure 6C**). However, blebbistatin treatment inhibited alignment of protrusions to the engineered fibers, particularly in HFF cells, were the distribution of angles to the fibers was flattened upon inhibition of contractility (**Figure 6D**). Expression of paxillin by HFF cells in aligned matrices remained diffuse upon treatment with blebbistatin (**Supplementary Figure 10**). The phenomenon of decreased protrusion alignment to the fibers was observed to a somewhat lesser extent in U87 cells (**Figure 6E**).

**Figure 6.**
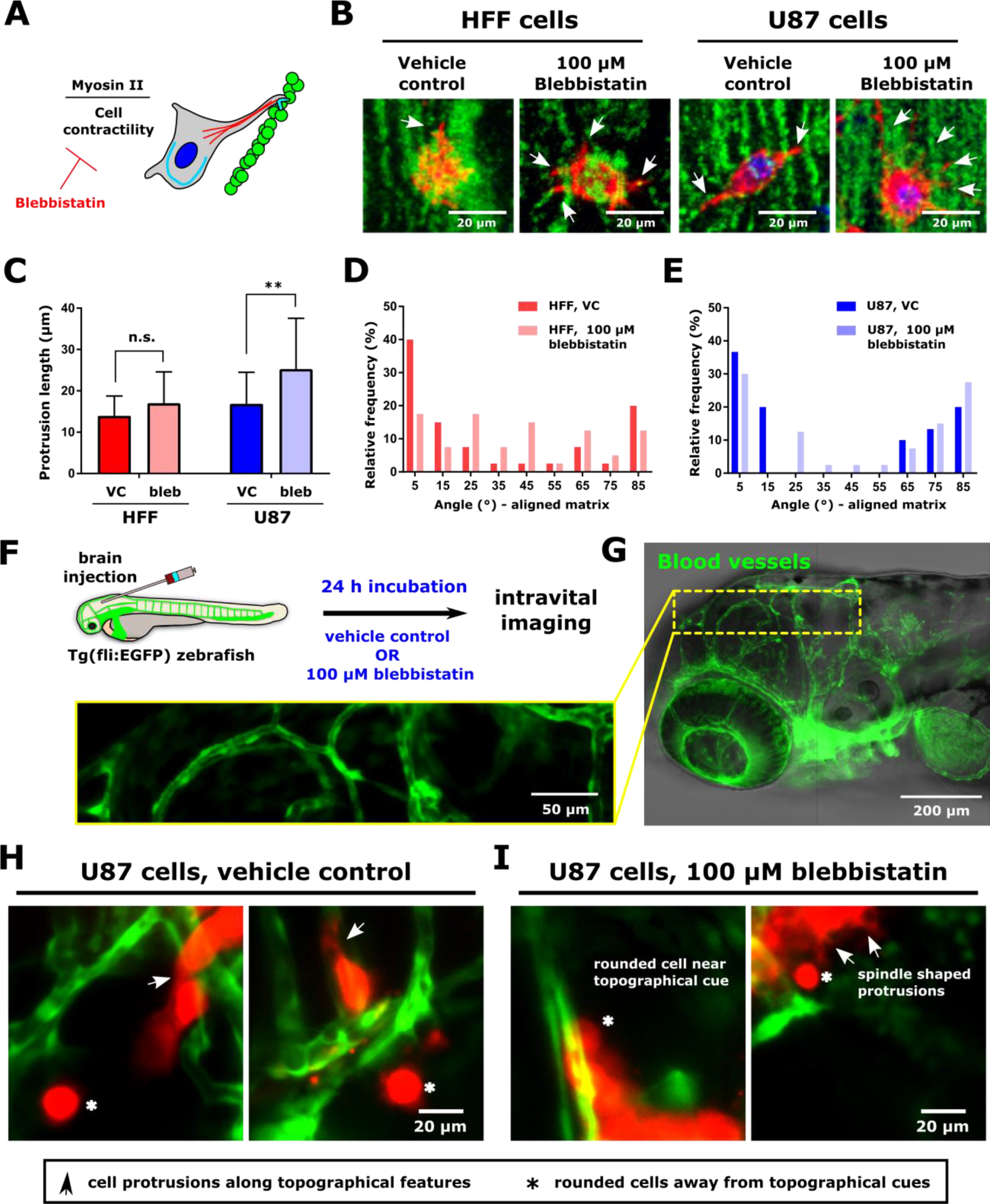
Modulation of protrusion alignment via inhibition of cell contractility in vitro and in vivo. (A) Schematic of contractility inhibition experiments. Myosin II-mediated cell contractility was inhibited with Blebbistatin. (B) Representative images of HFF and U87 cells in Matrigel matrices containing aligned fibronectin-containing particles upon treatment with 100 μM Blebbistatin or vehicle control (VC; DMSO). Arrows indicate cell protrusions. Particles are displayed in green, F-actin is displayed in red, and the nucleus is displayed in blue. Scale is indicated. (C) Cell protrusion length (mean ± standard deviation) in cells treated with 100 μM Blebbistatin or VC. (D) Distribution of angles between cell protrusions and the nearest fiber for HFF cells treated with 100 μM Blebbistatin or VC. (E) Distribution of angles between cell protrusions and the nearest fiber for U87 cells treated with 100 μM Blebbistatin or VC. In panels (C-E), either two matrices were analyzed per cell type and treatment. This resulted in analysis of 40 protrusions per condition for HFF cells and 30 (VC) or 40 (100 μM Blebbistatin) protrusions analyzed for U87 cells. (F) Schematic of in vivo experimental design. Cells were injected to the hindbrain of transgenic Tg(fli:EGFP) zebrafish, in which vascular epithelial cells express EGFP. After 24 h incubation fish water supplemented with 100 μM blebbistatin or VC, cells were imaged. (G) Overview image of zebrafish brain. Inset shows higher resolution of vessels in the brain. Vessels are displayed in green. Scale is indicated. (H) Images of U87 cells in the zebrafish brain following incubation in water supplemented with vehicle control. (I) Images of U87 cells in the zebrafish brain following incubation in water supplemented with 100 μM blebbistatin prior to imaging. Images are average intensity projections of confocal z stacks. Cells are displayed in red, and zebrafish blood vessels in red. Arrows indicate cell protrusions, while asterisks indicate rounded cells.

Aligned structures in vivo span varying lengths, from ECM fibers to guidance cues received from co-opting blood vessels. One question concerns whether the thickness of the fibers as well as the alignment is important in driving cell response. In our system, the fibrils were ~1-2 μm in thickness but show persistence of several microns. Thus, to begin assessing in vivo relevance, we employed an animal model wherein cell response to a topographical cue was readily observable via intravital microscopy. We injected HFF and U87 cells to the hindbrain of 2 days post fertilization (2 dpf) larval Tg(fli:EGFP) zebrafish, which are largely transparent and in which vascular endothelial cells express EGFP (**Figure 6F**). At 2 dpf, vessels in the zebrafish brain are wider than the fibers formed in the in vitro system (typically ~10-20 μm in width), with several regions in each fish where they were linear on the scale of a cell body length (**Figure 6G**).

We injected cells stained with a membrane dye to the zebrafish hindbrain and incubated the fish and cells for 24 hours post-injection in water supplemented with 100 μM Blebbistatin or the appropriate vehicle control (**Figure 6F**). To assess cell spreading in response to topographical cues, we imaged cells in the vicinity of blood vessels. Under control conditions, both HFF and U87 cells proximal to topographical cues elongated and extended protrusions, usually parallel to the long axis of the vessels (arrows, **Figure 6H** and **Supplementary Figure 11**). While protrusions were typically parallel to the vessels, some cells elongated perpendicular to these cues (**Figure 6H**, left panel), similar to our observations in the in vitro system. In contrast, cells away from vessels remained rounded (**Figure 6H**, asterisks, and **Supplementary Figure 11**). To assess whether inhibiting contractility would affect alignment in vivo in a similar manner to that observed in the engineered in vitro system, we treated a subset of U87 cells with 100 μM Blebbistatin overnight. Blebbistatin-treated U87 cells remained rounded along blood vessels (**Figure 6I**, left panel) or sent out thin, spindle-shaped protrusions in the absence of topographical cues, similar to what was observed in vitro (**Figure 6I**, right panel).

## DISCUSSION

The ECM provides important chemical signals within native tissue [2, 19]. In addition, physical properties of the microenvironment, such as tissue mechanics and surface topography, have been shown to modulate gene expression [2, 19]. In the model described here, cells receive a myriad of physical cues, namely microscale heterogeneities in rigidity, topographical cues due to the presence of aligned fibrils, and chemical signals from the matrix and assembled fibers. Using active microrheology, we quantified the mechanical variations present within a few micrometers of assembled fibers in the Matrigel matrix that was not resolved at the mm length scale using bulk rheology. We then used this system to characterize how cells respond to well-defined, introduced anisotropy in 3D for different types of ECM proteins. In this system, we assessed the first point of contact that cells make with the ECM, that is, formation of protrusions after embedding in a 3D environment. We used pharmacological and genetic perturbations of key proteins regulating protrusion dynamics and cell contractility to gain insights into the mechanisms governing this cell response. We determined that perturbation of myosin II abrogated the cells’ ability to “sense” the topographical cues.

Microscale mechanics are dependent upon local mesh geometry [43]. One concern is that the magnetic particles are rigid and thus will introduce local heterogeneities on the micrometer scale that may influence cell behavior. Compared to beads measured in unaligned gels, beads within one micrometer of the nearest fiber in aligned gels were significantly more rigid, whereas rigidity decreased with distance at 2-4 μm and was lower than in unaligned gels, possibly owing to changes in local concentration in the background matrix due to interactions with the nanochain fibers. Thus, one reason for the observed microrheological differences may be that the mesh size in aligned hydrogels differs from that in unaligned hydrogels. However, in our previous work [17], FRAP measurements using 150 kDa dextran indicated no differences in recovery in unaligned and aligned hydrogels compared to Matrigel alone. We confirmed these findings using a lower molecular weight dextran (10 kDa) in this work. The stiffness gradients observed in this work may contribute to protrusion generation or stabilization. Previous work has demonstrated that fibroblasts generate and retract protrusions to sense the underlying substrate stiffness [25], and that cell-scale increases in matrix stiffness near bundled collagen fibrils contribute to increased stability of cell adhesions to promote adhesion maturation in response to matrix alignment [44]. The increased matrix elasticity near the engineered fibrils presented here may direct or stabilize protrusions parallel to the fibers in a similar fashion. Protrusions directed perpendicular to the fibers are potentially initially directed along the nanogrooves between paramagnetic particles. Following protrusion generation, cell contractility may further deform the matrix to locally align protrusions and assembled fibrils.

In addition to differences in the magnitude of the complex modulus, we observed a difference in frequency dependence between the aligned gels and unaligned gels. Power law dependence of the complex modulus on frequency has been previously observed in a number of biomaterials, and various models have been proposed to explain this behavior according to the underlying dynamics of the constituent polymers (47-50). In fibrillar collagen gels polymerized at 2 mg/ml or 6 mg/ml at 4°C or 37°C, we previously observed power law exponents ranging from 0.66 – 0.74 [33], in the range predicted for a semi-flexible polymer network. Here, we observed exponents of ~0.6 in unaligned gels but only ~0.25 near fibers in aligned gels. Crossover frequencies at which G’’ exceeds G’ correspond to characteristic relaxation times and reflect the intrinsic time scale of energy dissipation processes inside the material. In the previously measured collagen gels, crossovers occurred at 300 Hz for the gels with fine mesh of small fibers and 2750 Hz for the gels with larger mesh and thicker, longer fibers [33]. Here, we observed unaligned gel crossovers at 1kHz and aligned gel crossovers at 10 kHz. At frequencies greater than 500 Hz, few physiological processes such as ion channel gating have been documented that might be influenced by external cues in this dynamic range [45]. Nevertheless, it remains to be seen how such a transition will modulate cell phenotypes.

At the level of matrix topography, we previously determined that the viscosity of the gel, concentration of the colloid particle, and duration of the applied magnetic field regulated the size and length of the fibers and interfiber spacing [17]. Here, using an alignment time of 15 minutes and a Matrigel 3D matrix, we obtained fibers of width ~1-2 μm, somewhat thinner than those observed in human dermis, which contains thick, long collagen bundles ranging from ~20-50 μm in diameter surrounded by a mesh of finer collagen fibers [46]. Overall, matrix organization was similar to that observed in human lung tumor slices, where stromal protein density and organization are variable (and includes regions with nearly parallel fibers), and gap sizes between fibronectin and collagen fibers range from ~5-15 um [47], and in lymphocyte-rich areas (for example, in the lymph nodes) in humans and mice, which contain thin fibronectin fibers with average spacing of ~15 μm and gaps ranging from 5 to >30 μm [47, 48]. Thus, cells were presented with three-dimensional topographies similar to those in ECM matrices *in vivo*. Aligned topographies are well-known drivers of cell polarization [49, 50].

Differences in the ECM proteins surrounding cells can also influence phenotype. Both normal and tumor cells secrete copious amounts of ECM proteins, and an overabundance of several ECM proteins is associated with abnormal tissue pathology [2]. In the case of cancer, tumor-conditioned stromal cells at both sites of primary tumors and metastases secrete copious amounts of proteins. Specifically, astrocytes secrete tenascin C within the brain, whereas fibroblasts secrete fibronectin in response to tumor derived cytokines [51–53]. Therefore, physiologically relevant biomaterials models would ideally incorporate both human cells and human matrix proteins, which we include by conjugating different ECM proteins to the magnetic particles forming the aligned fibers. For both HFF and U87 cells embedded in Matrigel matrices with aligned fibers, protrusion lengths increased and protrusion angles became oriented relative to the fibers in comparison to unaligned matrices, independent of the chemistry of the particles. Increased protrusion lengths in aligned matrices with BSA-conjugated particles compared to those in unaligned matrices and hydrogels devoid of particles suggest that protrusions in aligned matrices were not simply the result of higher local ECM protein density from the assembled fibers. Additionally, these results suggest that tumor cells may also use a similar mechanism to respond to topography as normal mesenchymal cells. However, we cannot rule out that cells behaved similarly simply because they may be using the same receptor to bind to these proteins. Moreover, cellular production of adhesion proteins could also contribute to generation of cell protrusions. We also note that the characteristics of the underlying matrix, for example, the ability of cells to break down non-porous matrices, effects cell behavior. The flexibility of the matrix engineering process opens the door for incorporating aligned fibers within chemically tunable 3D matrices to incorporate both topographical cues from alignment and mechanical and chemical properties characteristic of a given organ microenvironment.

Mechanistically, integrins act as the transmembrane anchor between the cell and the ECM, and other studies have elegantly elucidated role of adhesion-scale ECM presentation and integrin binding [54]. Heterodimers of an alpha and beta integrin subunits are used by cells for specificity of ECM binding. FN, tenascin C, and laminin are known to be ligands for β1 integrin with alpha subunits of alpha5, alpha6, and alpha9, respectively [22, 51, 53]. A migrating cell responds to local mechano-chemical cues by polarizing the membrane, with protrusions such as filopodia and lamellipodia at the leading edge [55]. Integrin β subunits are transported in the leading edge and facilitate binding with the ECM, which then drives a cascade of signals that may result in motility or proliferation. Integrin β1 has also been implicated as a key protein used by cells in contact guidance [26] and in protrusion generation to aligned collagen fibers [40]. Similarly, cells cultured on 2D grooves comparable in size to individual focal adhesions polarize along nanogrooves in a β1-integrin dependent manner, whereas fibroblasts cultured on nanocolumns show an increase in the number of filopodia [56, 57]. Thus, we interrogated the role of β1 integrin in regulating the observed cellular behavior and found that knocking down integrin β1 reduced the of generated protrusions in aligned Matrigel matrices. However, these protrusions coaligned with the engineered fibrils. While the presence of β1 integrin was necessary for protrusion generation, we did not observe large, mature focal adhesions in protrusions oriented parallel or perpendicular to the engineered fibrils in our system. This diffuse paxillin staining in response to uniaxial alignment cues is reminiscent of that recently observed in MDA-MB-468 cells on microcontact printed collagen lines [58]. Additionally, the size of paxillin-containing focal adhesions decreases for fibroblasts plated on viscoelastic vs. elastic 2D substrates, possibly due to energy dissipation due to the viscosity of the microenvironment and independent of the ECM protein presented on the surface [59]. This finding suggests that the viscoelasticity of the 3D matrix used in our studies may contribute to a loss of large focal adhesion plaques, even in the presence of alignment cues.

Because fascin is the main actin-bundling protein in filopodia, we reasoned that it may also be involved in protrusion generation in response to the aligned fibers within the Matrigel hydrogels. Using an inhibitor that specifically blocks fascin-dependent filopodial formation, we observed that the length of protrusions was reduced, while the ability to be aligned to the fibrillar structures remained intact. Fascin inhibition in *in vitro* and *in vivo* migration and metastasis assays show a reduction of cell migration and metastasis, and overexpression of fascin in 3D aggregates of mouse mammary epithelial cells cultured 3D collagen gels show a modest ability to sense aligned collagen fibers [60, 61]. Reductions in metastasis upon inhibition of fascin *in vivo* may therefore be partially due to decreased sensitivity to topographical cues, which have been implicated in metastatic progression [62].

Myosin II can affect the net rate of cellular protrusions such as lamellipodia and lamellae [63]. Inhibition with blebbistatin eradicates large actin bundles in the lamellum but the lamellipodium remains intact for cells cultured in 2D [63]. It further inhibits coalescence of actin into proto-bundles at the lamellipodium–lamellum interface which in turn increases protrusions [63]. Previous studies on 2D patterned substrates revealed that area and orientation of lamellipodial protrusions contribute to contact guidance [64]. In our 3D system comprised of fibronectin-conjugated particles aligned in a Matrigel hydrogel, the length of the protrusions following blebbistatin treatment remained unchanged for human fibroblasts and increased in U87 glioblastoma cells for the given concentration. Moreover, Myosin II mediates local cortical tension to guide endothelial cell branching morphogenesis and migration in 3D [65]. In the case of the human fibroblasts, the protrusions were randomly oriented. On the other hand, for the case of the GBM cells, even though the distribution of aligned protrusions is less robust than in the control case, there is still a fraction of protrusions that are aligned to the external fibers in our 3D cultures. GBM cells have been shown to be extremely plastic, and a given population is highly heterogeneous [66]. Thus, there may be cells within the population that are not as dependent on myosin II-mediated contractility. Taken together, these results suggest that understanding tissue heterogeneities may be important in modulating the response to both topographical and micromechanical cues.

Finally, cellular guidance and topographical cues in vivo are diverse and include fibrillar structures, such as collagen networks in tendons and the dermal layer in the skin, dense ECM networks, such as the basement membrane providing support to epithelial organs, and blood vessels [50, 67]. Vascular co-option is one strategy used by tumor cells to access nutrients in order to proliferate at distant sites [68, 69], and in cases of metastasis to the brain and breast, melanoma and lung cancer cells have been found to interact with the undulating basement membrane along the curvature of the existing blood vessels [68, 69]. Hence, cells respond to topographical cues for both thick and thin structures, and we examined whether cellular response to aligned fibrils in vitro could be extended to an in vivo system. We focused on U87 glioblastoma cells injected to the zebrafish hindbrain. Glioblastomas are highly migratory, where migration on brain vasculature facilitate widespread dissemination in vivo [70], and the mammalian and zebrafish brain share a number of similarities [71]. Upon injection, U87 cells in proximity to topographical cues from blood vessels extended protrusions parallel or perpendicular to the vessels. In contrast, blebbistatin treatment was sufficient to reduce alignment with blood vessels, and U87 cells also showed random protrusions away from blood vessels. Within the brain, glioma cells migrate like nontransformed, neural progenitor cells, extending a prominent leading cytoplasmic process followed by a burst of forward movement by the cell body that requires myosin II [66]. Thus, myosin II-regulated cell contractility may also be important in topographical sensing.

## CONCLUSIONS

We aimed to understand how cells respond to defined topographical cues within a 3D viscoelastic hydrogel. First, we integrated human cells and ECM proteins into a 3D hydrogel system containing aligned fibrils, where both cell type and ECM proteins reflect homotypic interactions, and characterized the micron-scale mechanical cues that cells receive in the vicinity of the fibrils. Optical-trap based active microrheology measurements revealed a local stiffness gradient in the vicinity of aligned fibrils when those fibrils are formed in Matrigel. This increased elasticity proximal to fibrils was similar to that observed in fibrillar collagen gels [44]. Human fibroblast and glioblastoma cells seeded in aligned Matrigel matrices generated longer protrusions than cells in unaligned matrices, and these protrusions were predominantly parallel or perpendicular to the local fiber orientation, independently of the ECM protein coating of the paramagnetic beads aligned to generate the fibrils. We observed that paxillin, a marker of stable focal adhesions, was mainly diffuse, even for cells cultured in hydrogels with aligned fibers. The effects of micron-scale stiffness gradients on the generation of protrusions must be considered when using this system. However, we note that the aspect ratios of HFF and U87 cells were unchanged when seeded on substrates ranging in stiffness from 0.5 kPa to 64 kPa, a larger dynamic range than that observed in the vicinity of the fibers (**Supplementary Figure 8**). Protrusion generation in aligned Matrigel matrices was dependent on both β1 integrin expression and fascin activity. However, protrusion alignment was governed in part by cell contractility. Taken together, these results suggest that there is an interplay between cell contractility and sensing topographical cues. The biomimetic platform presented here can be further tuned to provide fundamental insights into aberrant cell mechanosensing in physiologically-relevant microenvironments.

## ACKNOWLEDGMENTS

This research was supported by the Intramural Research Program of the National Institutes of Health, the National Cancer Institute. We thank Daniel Blair and Xinran Zhang and the Institute for Soft Matter Synthesis and Metrology at Georgetown University for access to and assistance with bulk rheology measurements. We also thank Ken Yamada, NIDCR, NIH for the kind gift of HFF cell lines. We thank Jayne Stommel, NCI, NIH for the kind gift of U87 cell lines. A. Hruska received a NIH GSOAR Summer fellowship.

## COMPETING INTERESTS STATEMENT

The authors declare no competing interests.

## DATA AVAILABILITY

The raw data required to reproduce these findings are available from Kandice Tanner, Ph.D., 37 Convent Dr., Bethesda, MD 20852. The processed data required to reproduce these findings are available from Kandice Tanner, Ph.D., 37 Convent Dr., Bethesda, MD 20852.

## SUPPLEMENTARY DATA

**Supplementary Figure 1.**
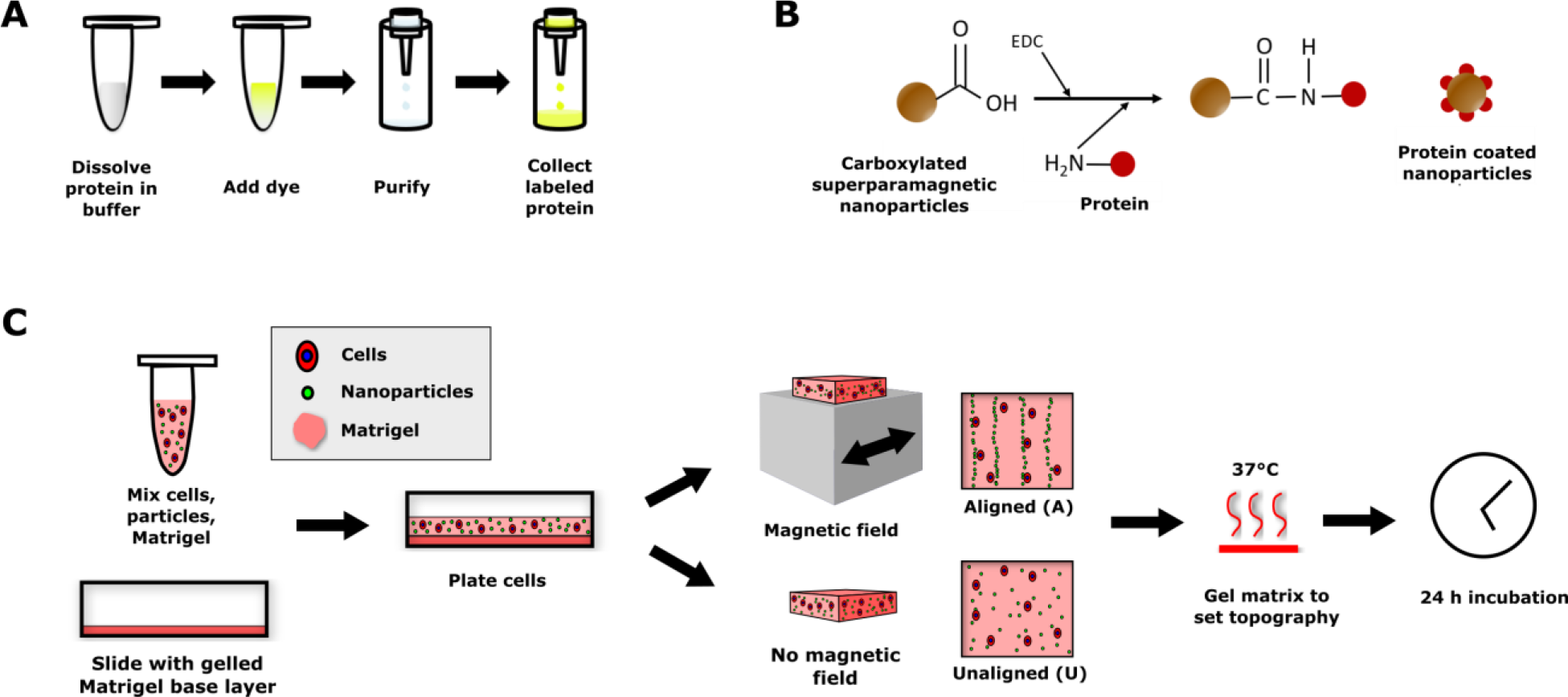
Detail of composite hydrogel fabrication process. (A) Schematic of procedure used to fluorescently label human extracellular matrix proteins. (B) Schematic representation of reaction scheme to conjugate human proteins with carboxylated superparamagnetic magnetic colloidal particles. (C) Schematic of 3D matrix assembly process. Human cells and superparamagnetic colloidal particles were suspended in Matrigel, plated on a glass slide containing a base layer of Matrigel, and either aligned in a magnetic field or left unaligned and dispersed throughout the matrix. Gels were then formed by heating at 37°C to set the matrix topography. Cells were fixed for analysis 24 h after seeding. Portions of this panel are repeated from Figure 1A to illustrate the entire matrix preparation process.

**Supplementary Figure 2.**
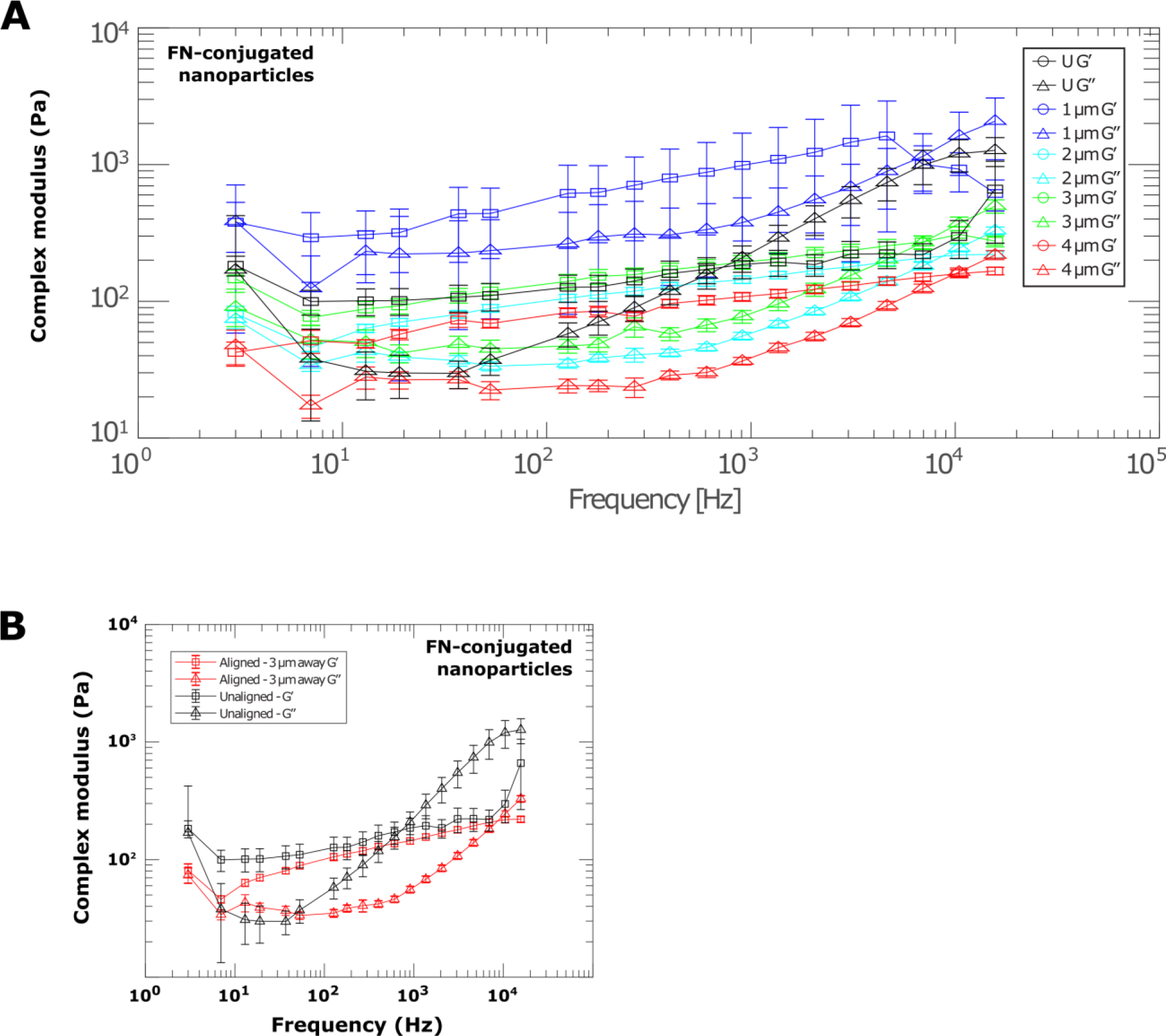
Frequency dependence of elastic and viscous components of complex modulus of aligned and unaligned gels via active microrheology. (A) Elastic (G’, circles) and viscous (G’’, triangles) components (mean ± SEM) of complex moduli of gels made with colloidal particles conjugated to human fibronectin. Moduli measured at beads in unaligned gels (black), or in aligned gels at distances of 1 μm (blue), 2 μm (cyan), 3 μm (green), or 4 μm (red) away from the nearest fiber. (B) Elastic (G’, squares) and viscous (G’’, triangles) components (mean ± SEM) of complex moduli of gels made with colloidal particles conjugated to human fibronectin. Moduli measured at beads in unaligned gels (black), or in aligned gels at distances of 3 μm away from the nearest fiber (red). Data is replotted from Supplementary Figure 2A for clarity. For all microrheology measurements, samples were measured in triplicate, with at least 30 beads per sample analyzed.

**Supplementary Figure 3.**
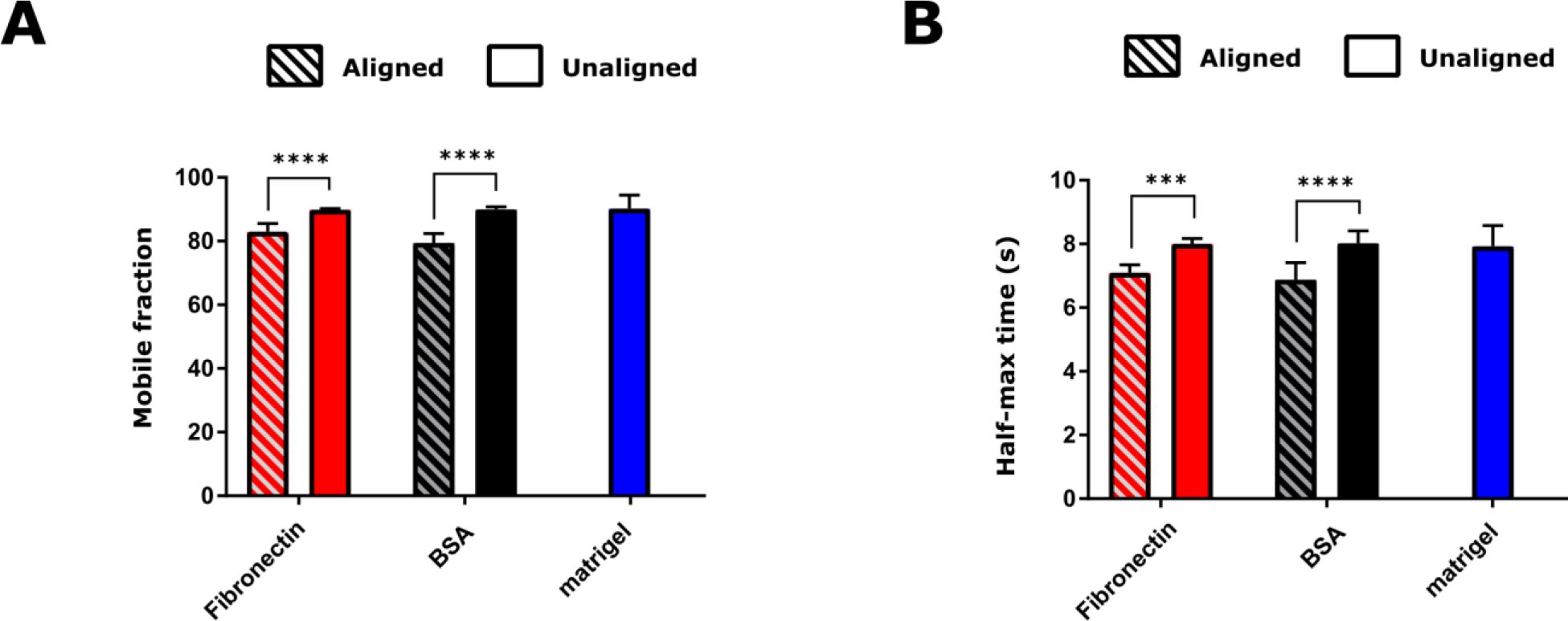
Fluorescence recovery after photobleaching in aligned and unaligned Matrigel matrices. (A) Mobile fraction and (B) half-maximum time in aligned and unaligned Matrigel matrices containing colloidal particles conjugated to fibronectin or BSA, or in Matrigel matrices without added particles, from fluorescence recovery after photobleaching experiments. For each condition (particle protein coating and alignment status), three independent regions from two gels were measured. These measurements were grouped to obtain N=6 values prior to statistical comparisons. ***, p<0.01 and ****, p<0.0001 by Sidak’s multiple comparisons test following two-way ANOVA.

**Supplementary Figure 4.**
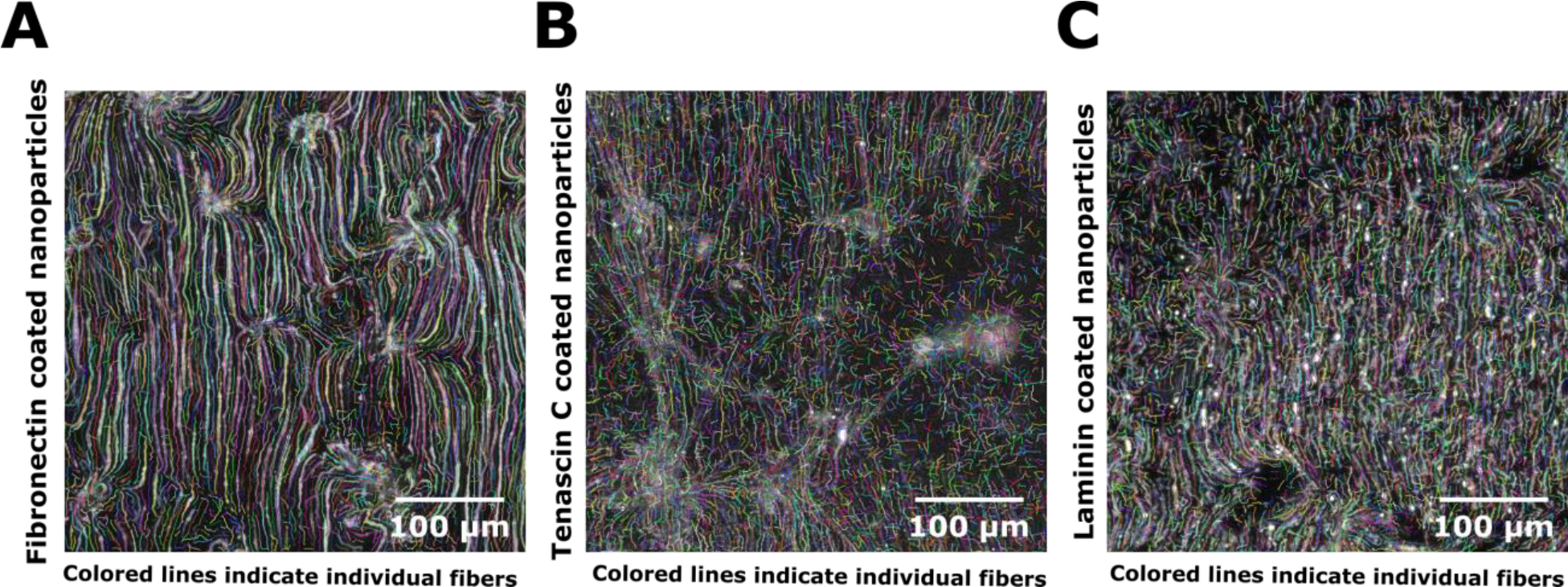
Analysis of fiber properties in aligned matrices containing protein-conjugated colloidal particles. Representative images of fibers formed of (A) fibronectin-conjugated colloidal particles, (B) tenascin C-conjugated colloidal particles, and (C) laminin-conjugated colloidal particles following segmentation using the ctFire fiber analysis toolbox.

**Supplementary Figure 5.**
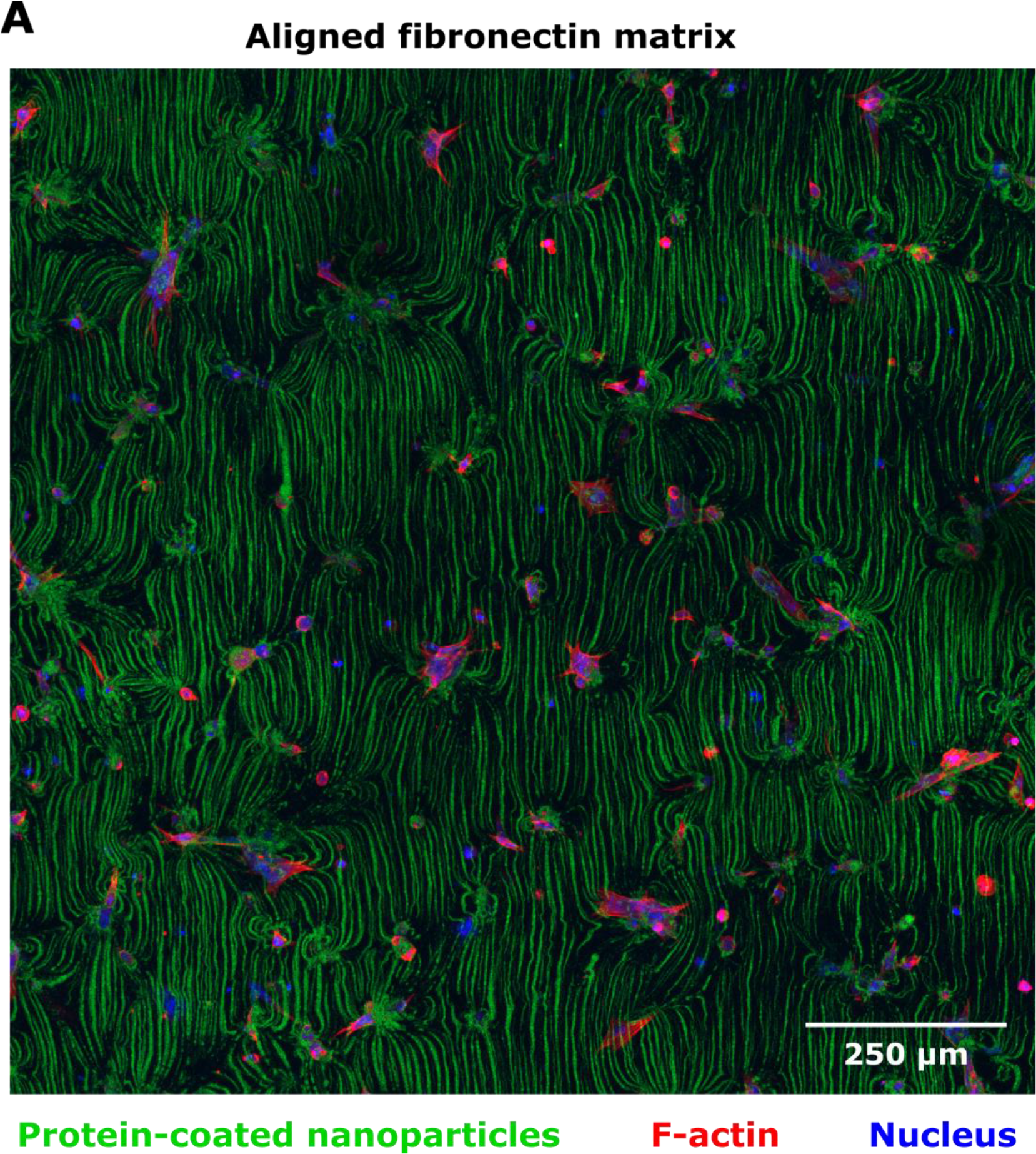
Large area, field-based fiber patterning. Representative images of HFF cells seeded in an aligned matrix with fibronectin-coated colloidal particles. Using the magnetic-field based alignment technique, large areas (>1 mm^2^) can be rapidly patterned around cells embedded in 3D matrices. Colloidal particles are displayed in green, F-actin in red, and the nucleus in blue. Image is maximum intensity projection of confocal tile scan slices. Scale bar = 250 μm.

**Supplementary Figure 6.**
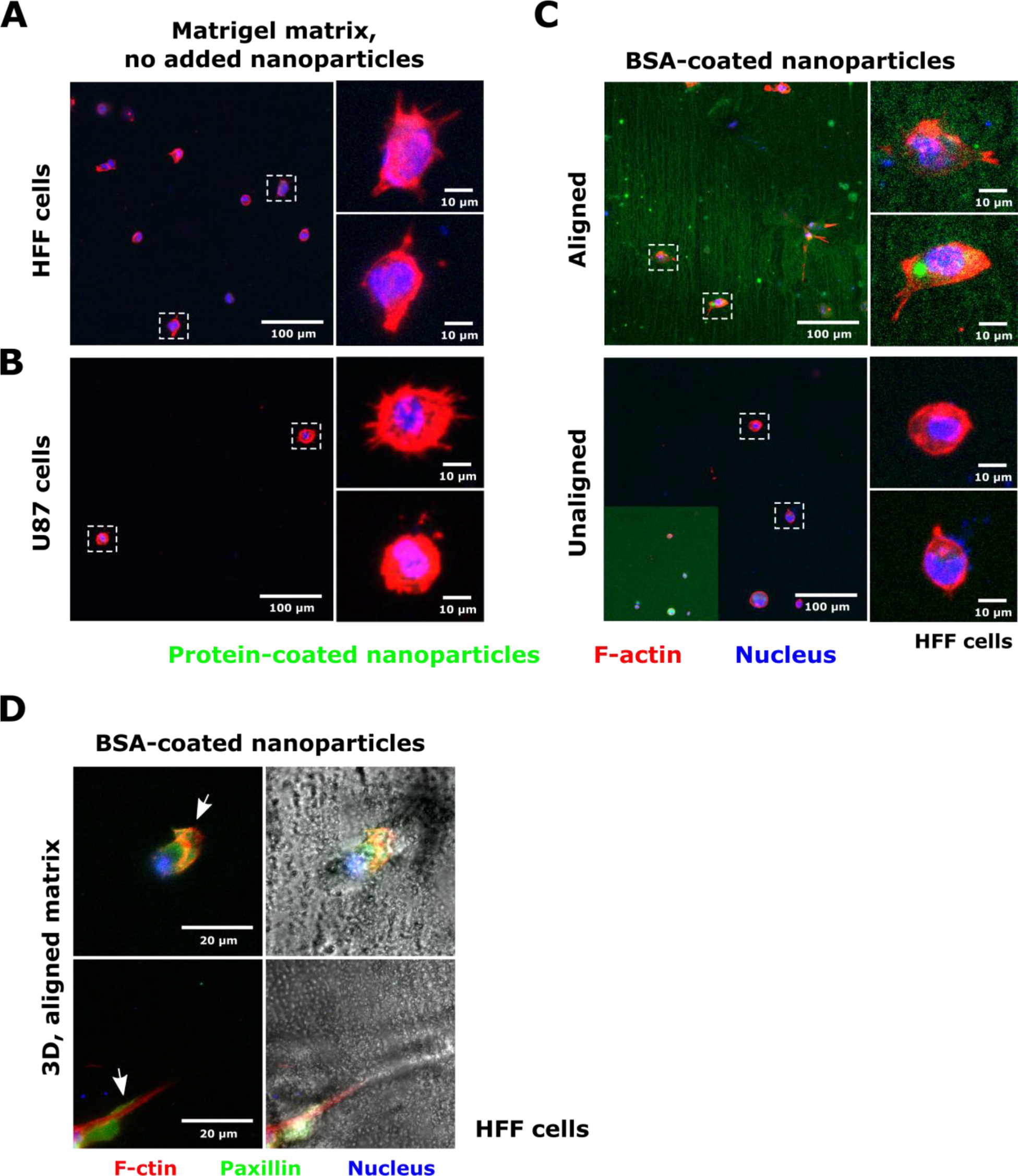
Cell morphology in Matrigel matrices without added colloidal particles and in gels containing BSA-conjugated colloidal particles. Representation images of (A) HFF and (B) U87 cells dispersed in a 3D Matrigel matrix no added Magnetic colloidal particles. (C) Representative images of HFF cells embedded in aligned and unaligned matrices containing colloidal particles conjugated to bovine serum albumin (BSA). Inset in panel C shows unaligned matrix with lookup table adjusted to show presence of dispersed fluorescent particles. Magnetic colloidal particles are displayed in green, F-actin is displayed in red, and the nucleus is displayed in blue. In each panel, overview images are shown, with insets to show detailed cell morphology (cell position in larger image indicated by dashed white boxes). Images are maximum intensity projections of confocal slices containing aligned fibers, or of cells embedded in 3D. Scales are indicated in each image. (D) Representative images of HFF cells in aligned Matrigel matrices containing BSA-conjugated nanoparticles and expressing a GFP-paxillin biosensor. Images are maximum intensity projections of confocal slices containing colloidal particles. F-actin is displayed in red, paxillin in green, and the nucleus in blue. Brightfield images are shown to illustrate particle alignment. Arrows indicate cell protrusions in the plane of the fibers. Scale is indicated.

**Supplementary Figure 7.**
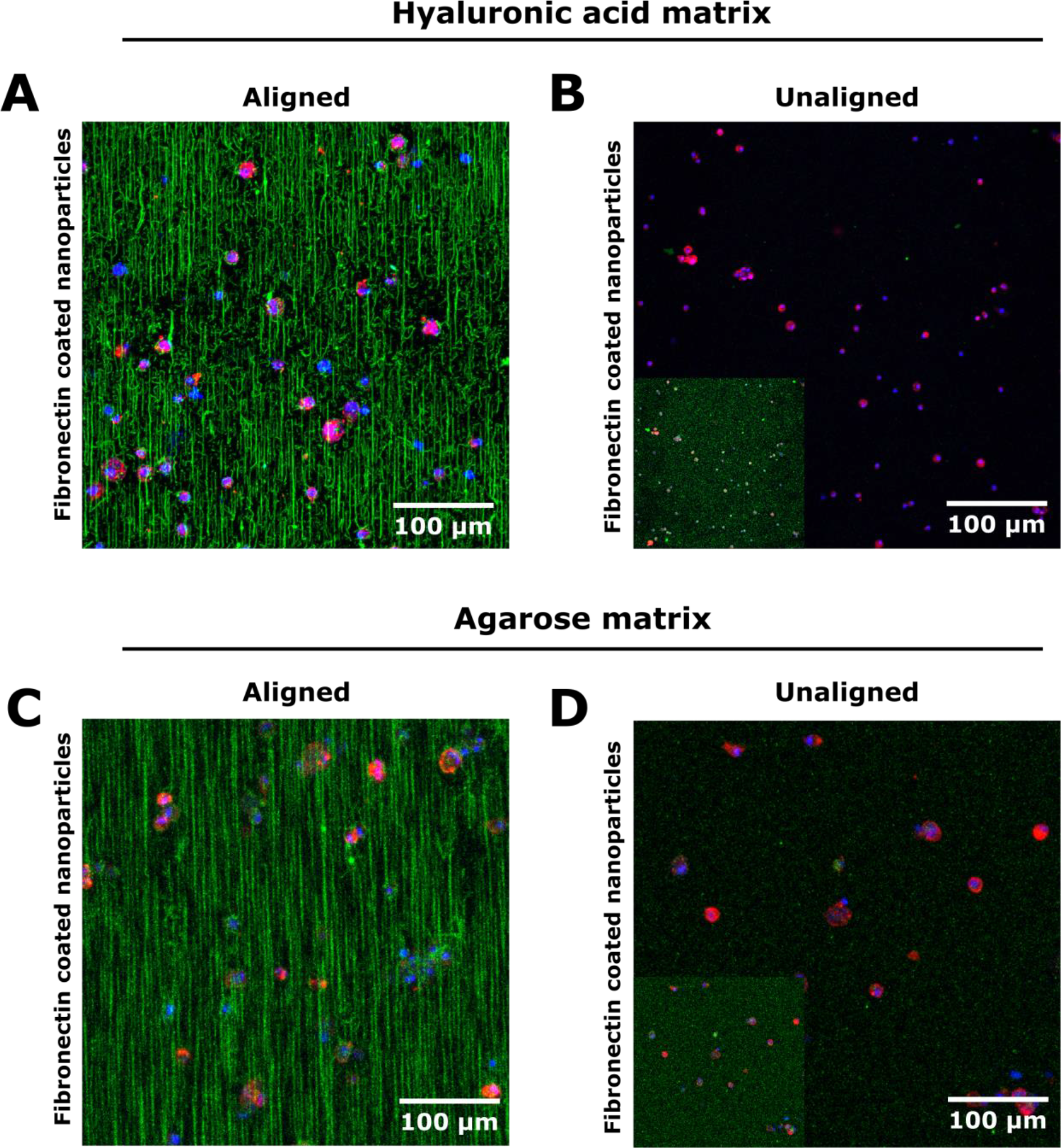
Engineered fibers in multiple 3D cell culture matrices. Human foreskin fibroblasts were embedded in hyaluronic acid matrices containing (A) aligned or (B) unaligned fibronectin-conjugated colloidal particles, or agarose matrices containing (C) aligned or (D) unaligned fibronectin-conjugated colloidal particles. Insets in unaligned matrix examples show the same unaligned matrix image with lookup table adjusted to show presence of dispersed fluorescent particles. Colloid particles are displayed in green, F-actin in red, and the nucleus in blue. Images are maximum intensity projections of confocal slices containing aligned fibers. Scales are indicated.

**Supplementary Figures 8.**
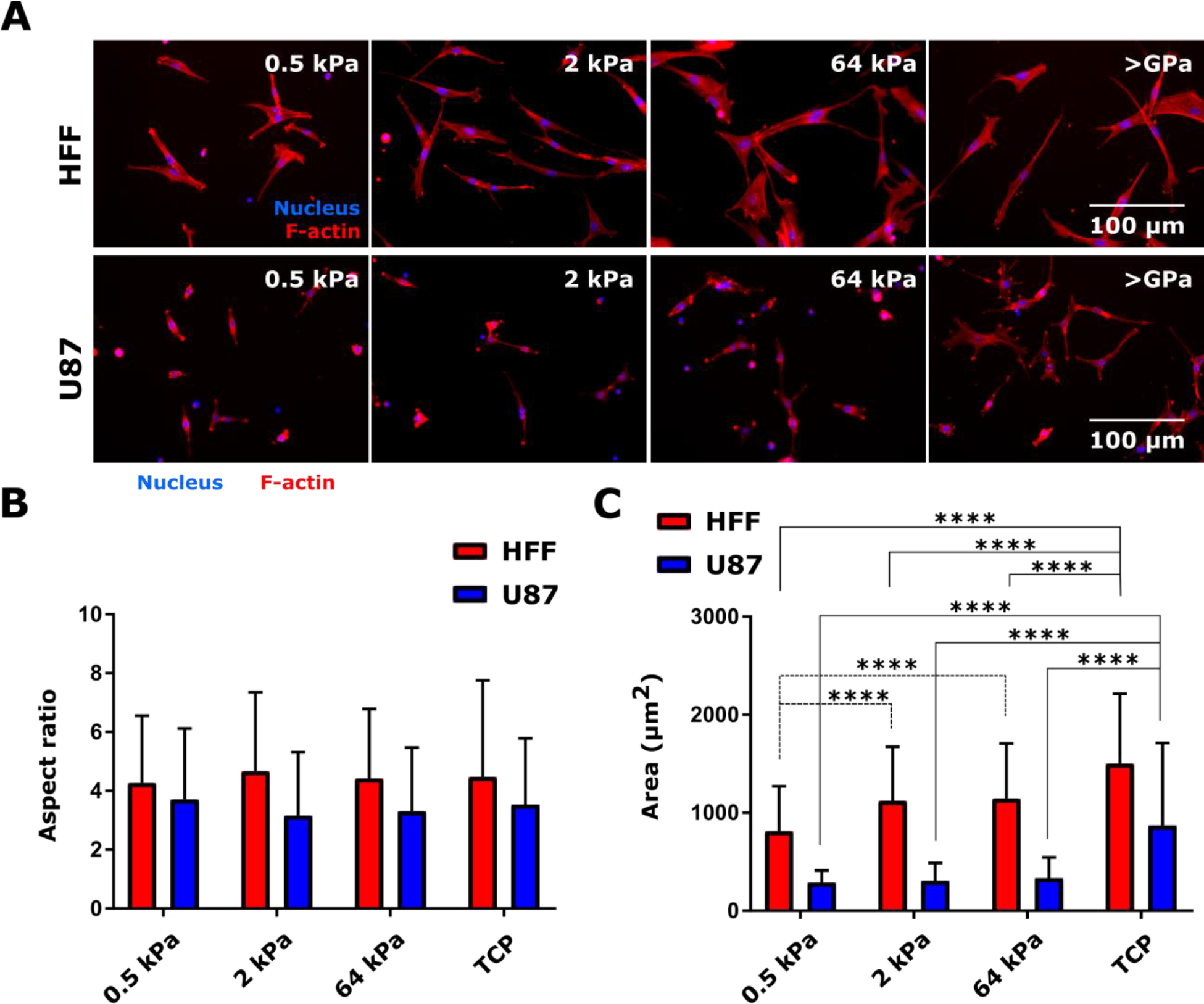
Effect of substrate stiffness on HFF and U87 cell morphology. (A) Representative fluorescence images of HFF and U87 cells seeded on silicone substrates with elastic modulus of 0.5, 2, or 64 kPa, or on tissue culture plastic (TCP, >GPa stiffness). Surfaces were coated with 1 μg/ml human fibronectin. F-actin is displayed in red, and the nucleus in blue. Scale is shown. (B) Aspect ratio (mean ± standard deviation) and projected cell area (mean ± standard deviation) as a function of cell type and substrate stiffness. Area and aspect ratio values among conditions were compared using two-way ANOVA with Tukey’s multiple comparisons post-test between all combinations of substrate stiffness for a given cell type. ****, p<0.0001. Measurements were made for two samples per cell type and substrate stiffness, and samples were prepared simultaneously. At least 73 cells were analyzed for each condition.

**Supplementary Figure 9.**
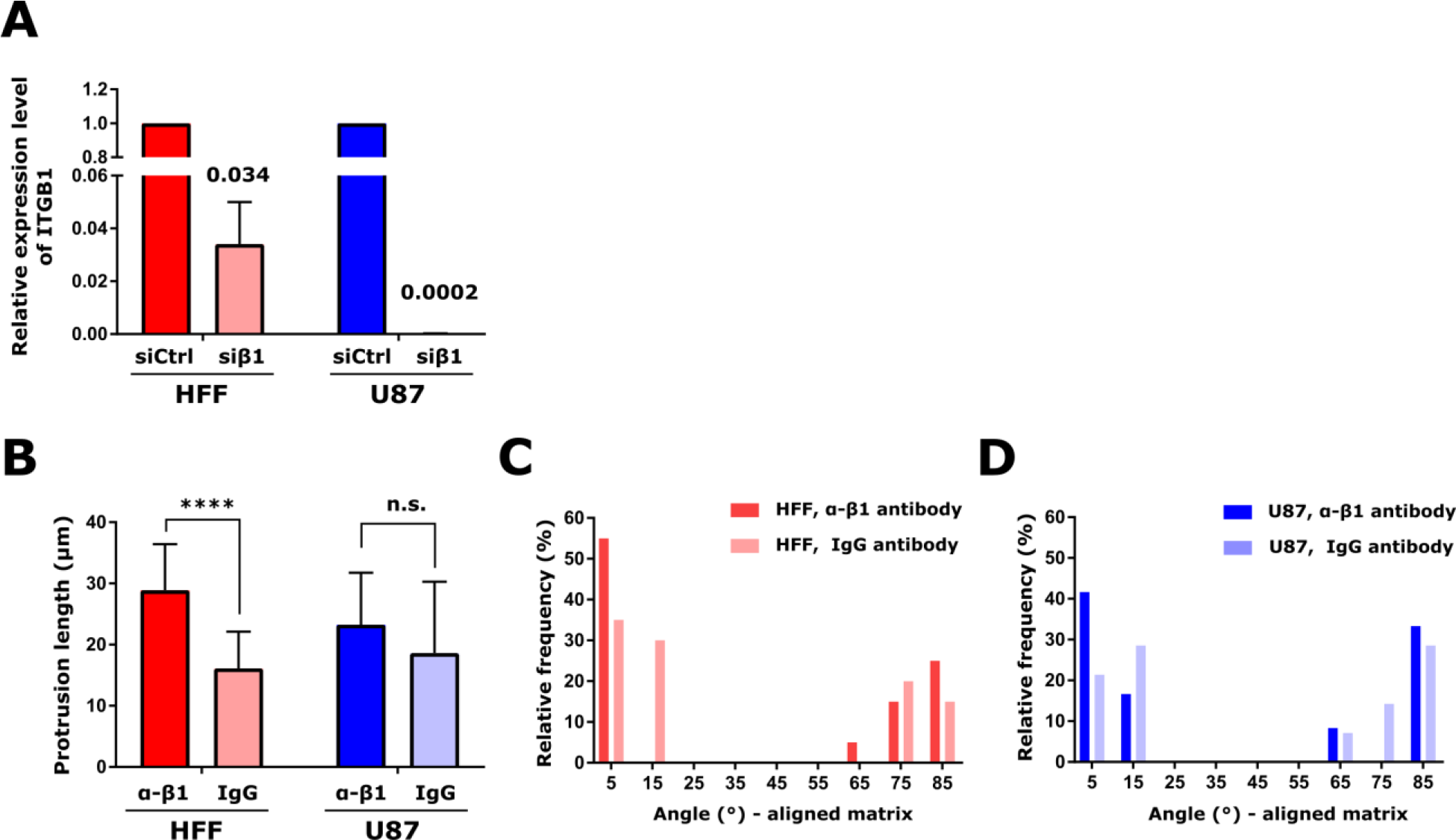
Knockdown efficiency and effects of integrin β1 function blocking on cell protrusion and protrusion angles in aligned matrices. (A) Relative expression of integrin β1 (ITGB1) in HFF or U87 cells transfected with non-targeting control siRNA or siRNA directed against integrin β1. Expression was assessed via qtPCR with relative expression calculated from gene ∆∆cT. GAPDH was used as the housekeeping gene. Numbers indicate relative expression upon knockdown. (B) Cell protrusion length (mean ± standard deviation) in cells treated with 30 μg/ml function-blocking antibody against integrin β1 (α-β1) control IgG antibody (IgG). (C) Distribution of angles between cell protrusions and the nearest fiber for HFF cells treated with 30 μg/ml function-blocking antibody against integrin β1 (α-β1) control IgG antibody (IgG). (D) Distribution of angles between cell protrusions and the nearest fiber for U87 cells treated with 30 μg/ml function-blocking antibody against integrin β1 (α-β1) control IgG antibody (IgG). In panels B-D, one matrix was analyzed per cell type and treatment, with 20 protrusions per condition analyzed for HFF and 12 and 14 protrusions analyzed for U87 α-β1 and IgG cells, respectively.

**Supplementary Figure 10.**
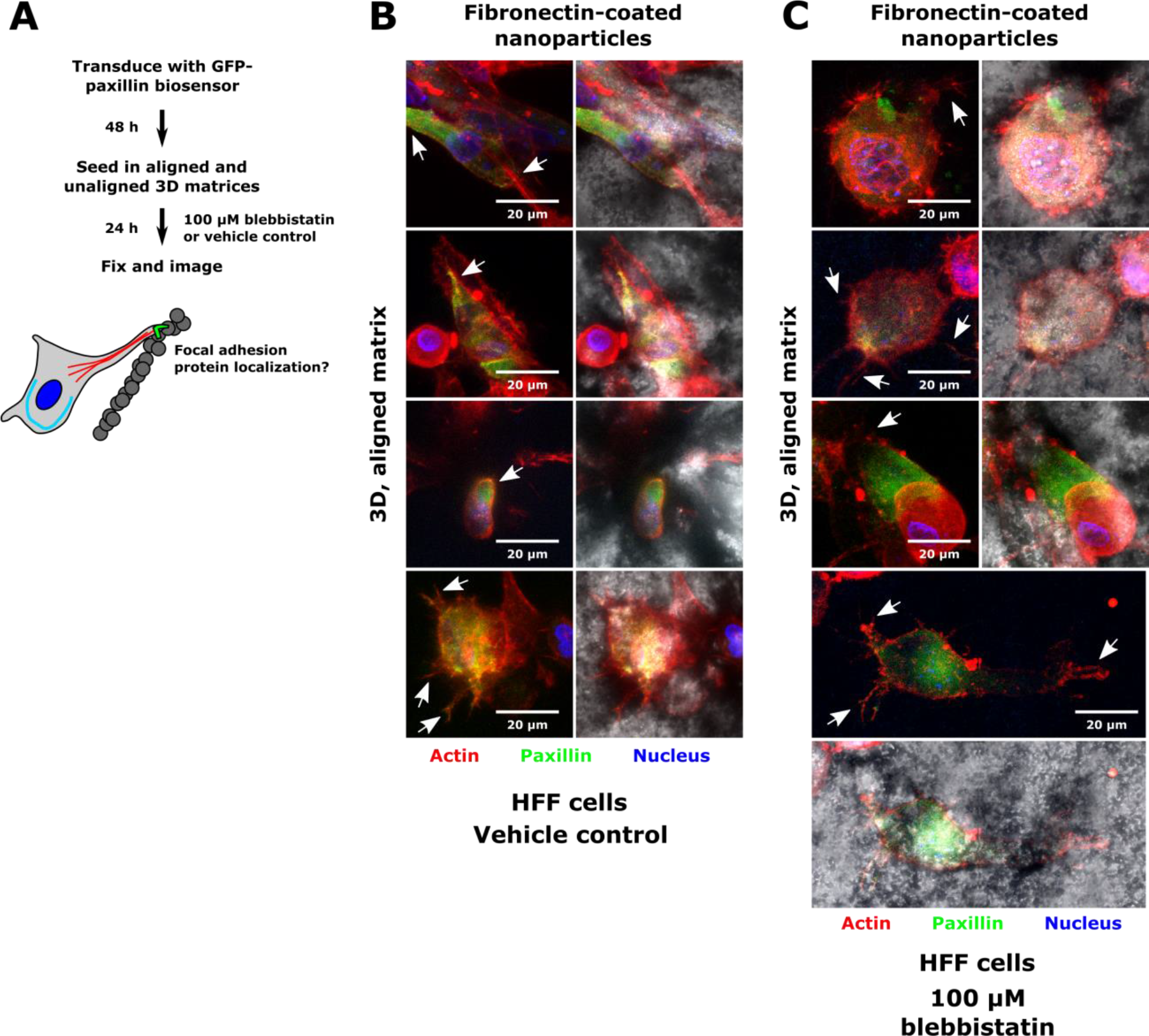
Human foreskin fibroblast (HFF) paxillin expression upon blebbistatin treatment. (A) Schematic of experiment to assess focal adhesion protein localization in HFF cells in 3D matrices in the presence of blebbistatin. Cells were transduced with a GFP-paxillin lentiviral biosensor prior to being embedded in aligned and unaligned Matrigel matrices in the presence of 100 μM blebbistatin or vehicle control. Cells were fixed, stained, and imaged after being embedded in matrices for 24 h. (B) Representative images of HFF cells in aligned Matrigel matrices containing fibronectin-conjugated nanoparticles and expressing a GFP-paxillin biosensor in the vehicle control case. (C) Representative images of HFF cells in aligned Matrigel matrices containing fibronectin-conjugated nanoparticles and expressing a GFP-paxillin biosensor after treatment overnight with 100 μM blebbistatin. In panels (B,C), images are maximum intensity projections of confocal slices containing colloidal particles. F-actin is displayed in red, paxillin in green, and the nucleus in blue. Brightfield images are shown to illustrate particle alignment. Arrows indicate cell protrusions in the plane of the fibers. Scale is indicated.

**Supplementary Figure 11.**
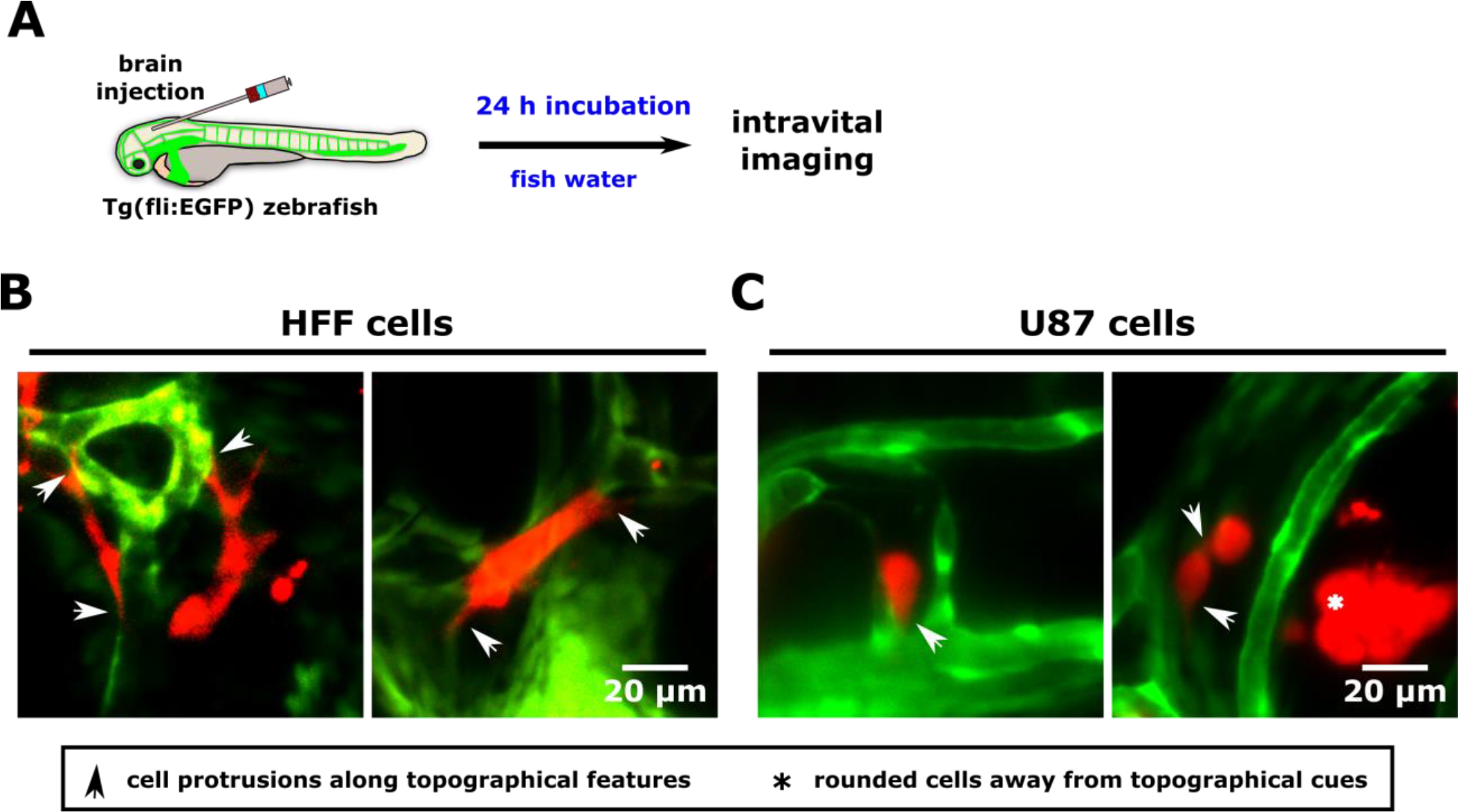
Alignment of cells to topographical cues in the in vivo zebrafish brain microenvironment. (A) Schematic of experimental design. Cells were injected to the hindbrain of transgenic Tg(fli:EGFP) zebrafish, in which vascular epithelial cells express EGFP. After 24 h incubation in fish water, cells were imaged. (B) Images of HFF cells in the zebrafish brain following incubation in fish water. (C) Images of U87 cells in the zebrafish brain following incubation in fish water. Images are average intensity projections of confocal z stacks. Cells are displayed in red, and zebrafish blood vessels in red. Arrows indicate cell protrusions, while asterisks indicate rounded cells.

**Supplementary Video 1.**
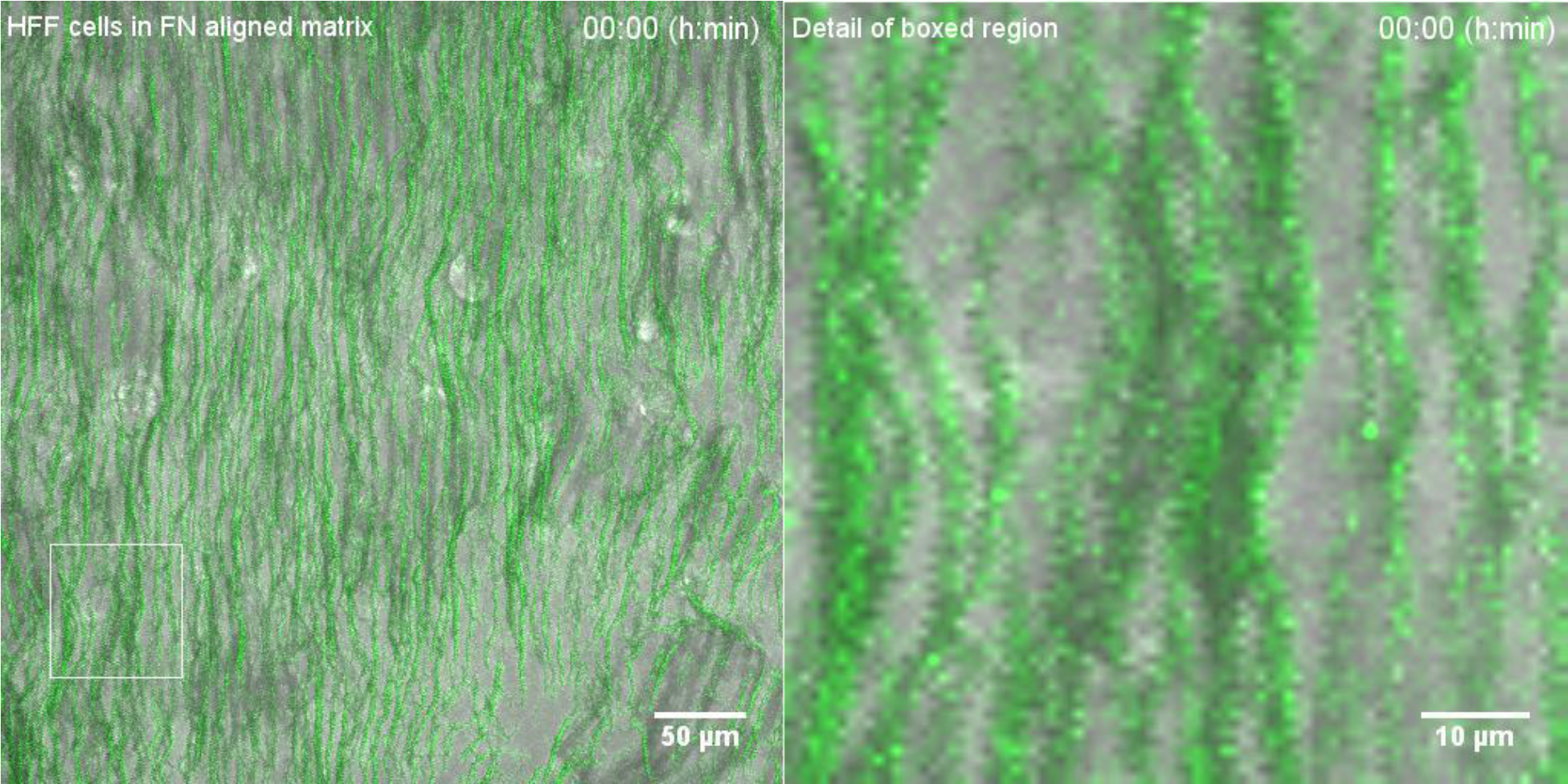
Time-lapse video of HFF cell protrusion and contraction in aligned matrix. HFF cells (bright field) were plated in aligned matrices containing fibronectin-conjugated colloidal particles (green). Time-lapse video shows cells protruding in and contracting the matrix. White box indicates region of interest shown in detail in right panel. Images show maximum intensity projection of confocal z slices and were acquired every 10 min, starting immediately after matrix topography was set.

**Supplementary Video 2.**
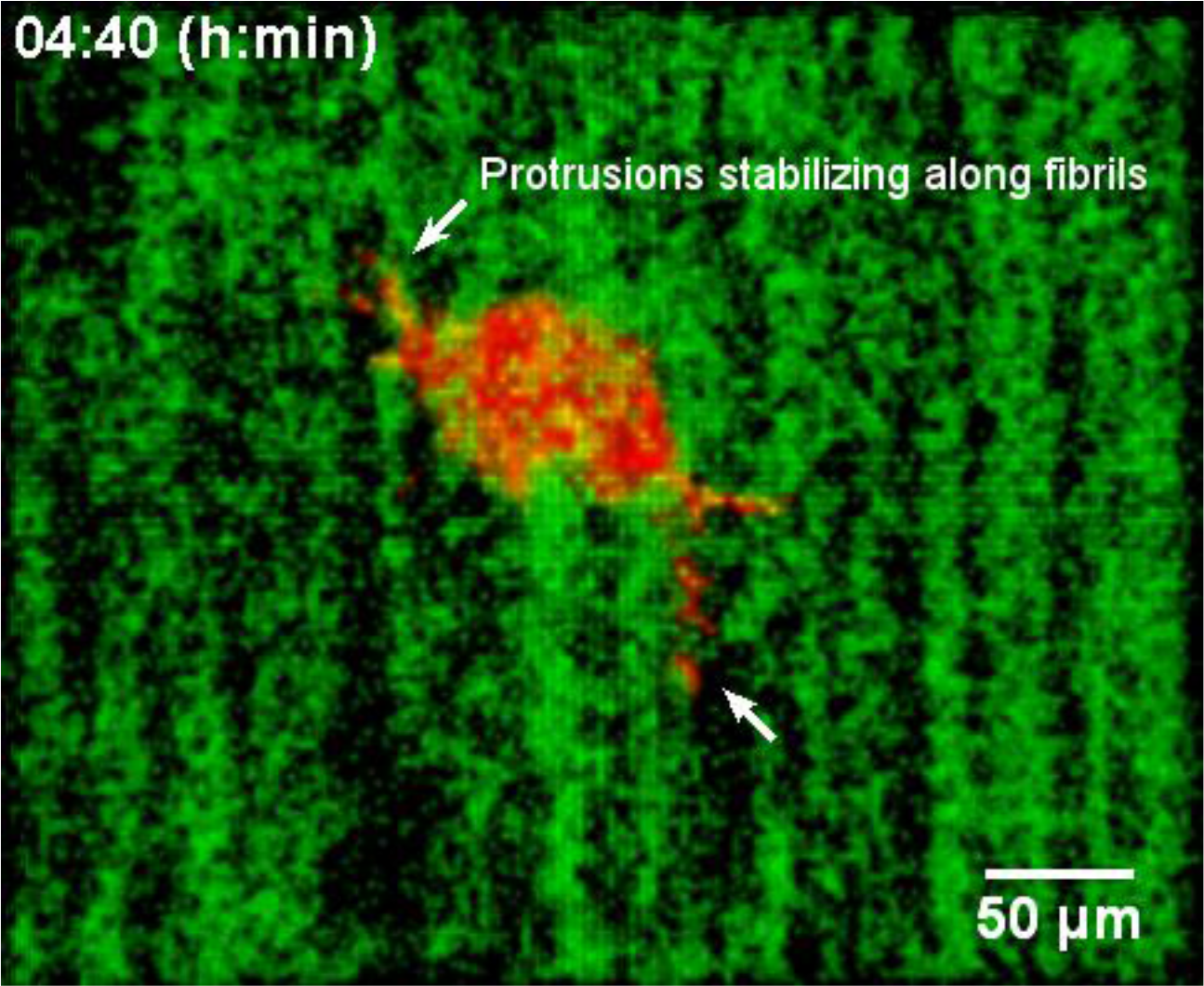
U87 cell actin cytoskeletal imaging reveals protrusion maturation along aligned fibers. A U87 cell was transduced with a LifeAct adenovirus (red) and plated in an aligned matrix containing fibronectin-conjugated colloidal particles (displayed in green). Video shows 3D reconstruction of confocal z slices, which were acquired every 10 min. Time stamps and scale bars are indicated. Arrows point to protrusions forming along fibrils.

